# Evolutionary divergence V-ATPase function in macropinocytic cup remodeling

**DOI:** 10.64898/2026.08.10.743417

**Authors:** Bhagyashree Chordiya, Navyaka Padavala, Amisha Sharma, Kadambini Pradhan, Kuldeep Verma

## Abstract

Macropinocytosis is an evolutionarily conserved process from unicellular eukaryotes to mammals. This process relies on actin-rich protrusions beneath the plasma membrane to form a macropinocytic cup. Traditionally, V ATPase has been best studied for its role in late endosomal acidification. However, the specific roles of V-ATPase interaction with cytoskeletal machinery and its crosstalk with lipids in the formation of macropinocytic cups have not been investigated. Here, we uncover an unprecedented role of V-ATPase in shaping the macropinocytic cup in the pathogenic amoeba *Entamoeba histolytica*. Our results showed that the *Entamoeba* V-ATPase complex is associated with F-actin at macropinocytic cups. Moreover, we found that the V-ATPase complex is dynamically recruited to the macropinocytic cup and dissociates from the macropinosome. We further demonstrated that the V-ATPase V1B subunit directly binds to actin and uniquely promotes actin polymerization. The pH biosensor phosphatidic acid, revealed abundant V-ATPase complex and cytoskeleton associated proteins in its interactome. Surprisingly, the V1B subunit showed reproducible binding to phosphatidic acid and this interaction modulates actin polymerization. Collectively, our work highlights the novel role of V-ATPase in directly driving actin polymerization, in conjunction with phosphatidic acid, to shape the macropinocytic cup.

## Introduction

Macropinocytosis is a form of non-selective endocytosis. This process is evolutionarily conserved from unicellular eukaryotes to mammals. It plays a crucial role in cellular physiology by facilitating nutrient uptake, immune responses, and cellular signaling [1]. This pathway is typically initiated in response to external or internal cues. Upon stimulation, the cell membrane extends outward to form ruffles through the coordinated action of lipids including phosphatidic acid (PA) and phosphoinositide 3 kinase (PI3K)-mediated actin polymerization [2, 3]. These ruffles curve back towards the cell surface, enclosing extracellular fluid and forming a cup-like structure [4]. The closure of this structure results in the formation of a nascent macropinosome, which traps the engulfed fluid. Following their formation, the actin coat surrounding the macropinosome dissociates, and the vesicle undergoes acidification through fusion with endo-lysosomal compartments, enabling the degradation and processing of the internalized material [5].

Nearly a billion years ago, macropinocytosis is thought to have arisen in the common ancestor of Amoebozoa and Opisthokonta, a lineage that includes animals, highlighting its deep evolutionary roots [6]. Within Amoebozoa, free-living members such as *Dictyostelium* species have become invaluable model organisms for studying macropinocytosis, owing to their robust and constitutive use of this pathway for nutrient uptake [7]. In contrast, certain amoebozoans, including *Acanthamoeba* and *Entamoeba* species, have adapted this process for pathogenic purposes, enabling them to invade and establish infections within human or animal hosts [8].

*Entamoeba histolytica* is a human pathogen and causative agent of amoebiasis. Transmission of this pathogen occurs not only via contaminated food and water but also through sexual contact, both homosexual and heterosexual [9]. These infections occur worldwide, however most cases are reported from India, Bangladesh, Africa, Mexico, and parts of Central and South America [10]. Each year, the disease affects approximately 50 million individuals and leads to around 100,000 deaths, disproportionately impacting children under five years and immunocompromised adults, including those living with HIV [11, 12]. Amoebiasis is the third parasitic disease causing death worldwide, after malaria and schistosomiasis. In *E. histolytica*, endocytic processes such as macropinocytosis, phagocytosis, and trogocytosis are indispensable for nutrient acquisition, tissue invasion, and immune evasion [13–15]. This parasite showed tremendous macropinocytic activity comparable to *Dictyostelium discoideum*, with both organisms capable of internalizing approximately 5–15% of their cell volume per hour under active conditions [16]. Although, macropinocytosis related studies on *Entamoeba* are still limited, existing literature indicate that this process is highly conserved with higher eukaryotes [17]. It relies on actin-driven process involves small GTPases, PI3-kinase signaling and actin regulators [18–20]. Recent findings suggest that the impaired intracellular pH of acidic compartments disrupts the trophozoite’s ability to perform several endocytic processes, highlighting the importance of lysosome-mediated functions in host cell killing [21]. Previously, it has been shown that the pharmacological inhibition of V-ATPase leads to impairment of phagosome acidification in amoebic trophozoites [22].

In most eukaryotic cells, the acidification of endolysosomal compartments is primarily regulated by the activity of vacuolar-type H⁺-ATPase (V-ATPase) pump [23, 24]. V-ATPase is a multisubunit proton pump that consists of two major domains, V1 and V0. The peripheral V1 domain contains eight distinct subunits (A-H), with a central catalytic hexamer formed by alternating A and B subunits that hydrolyzes ATP to generate mechanical energy [25]. This energy drives the rotation of a central stalk made up of subunits D and F, while three peripheral stalks composed of subunits E and G stabilize the hexamer and prevent its rotation [26]. On the other hand membrane-integrated V0 domain consists of five subunits (a, c, c’, c’’, and d), where subunit d connects the rotary stalk to a proteolipid c-ring embedded in the membrane [27]. This ring, composed of isoforms c, c′, and c″, rotates in response to the torque generated by ATP hydrolysis, enabling proton translocation across the membrane in coordination with subunit a, which facilitates the entry and exit of protons in the lumen [28]. However, the C subunit of V1 domain, along with the regulator of the H^+^- ATPase of vacuoles and endosomes (RAVE) complex, plays a regulatory role in the assembly of V-ATPase complex [29].

Beyond their canonical role in acidification, V-ATPase subunits particularly those in the V1 domain have emerged as regulators of cytoskeletal dynamics [23]. In multiple kingdoms, it has been shown that the V-ATPase V1B, V1C and V1E subunits interact with actin, influencing its organization [30, 31]. *Arabidopsis thaliana* V1B subunit contains N-terminal profilin-like motifs that directly interact with actin [32]. In *Manduca sexta* V1C subunit interacts with both G-actin and F-actin [33]. The ectopic expression of the *Drosophila melanogaster* V1C subunit increases the acidification of vacuoles, and induces a c-Jun N-terminal kinase (JNK)-dependent invasion [34]. In *Dictyostelium discoideum*, the V-ATPase associates with F-actin, and its removal from lysosomes is driven by actin polymerization, but the key nucleation-promoting factor involved is WASP and SCAR homologue (WASH) [35]. Further, a study showed that actin depolymerization promotes disassembly of the V-ATPase complex in HeLa cells [36]. Recently, it has been shown that the plasma membrane V-ATPase-mediated transport of cholesterol is required for macropinocytosis in mammalian cells [37]. However, the roles of V-ATPase in *Entamoeba* biology remains unclear. The parasite genome encodes all subunits required for the assembly of a V-ATPase complex, but the functional characterization of this complex is poorly explored [38, 39].

Nevertheless, the direct role of V-ATPase in actin remodelling during the early stages of macropinocytosis remains poorly understood. To address this question, we investigated the role of V ATPase in actin remodelling during early macropinocytosis in *E. histolytica*. We employed a multidisciplinary approach combining biochemical assays, cell biology techniques, high-resolution fixed and live-cell imaging, proteomics, and quantitative analysis. Our study demonstrates that dynamic V-ATPase complex recruitment occurs at the macropinocytic cup. Notably, the evolutionarily conserved V1B subunit directly binds with actin and uniquely regulates actin polymerization. Furthermore, our PA pulldown analysis revealed consistent binding of V-ATPase subunits particularly V1B, and this interaction fine-tune the actin polymerization. Taken together, these findings support a model in which V-ATPase coordinates macropinocytic cup formation through a regulatory axis involving actin polymerization and PA interaction.

## Results

### *Entamoeba* V-ATPase localizes on the plasma membrane and acidic compartments

To investigate the functional architecture of the V-ATPase complex in *Entamoeba*, we conducted an *in silico* analysis to identify its constituent subunits (Supplementary Information; Tables S1, S2). Subsequently, we selectively cloned and engineered constructs with a 3× hemagglutinin (HA) tag at the N-terminus of the V1A, V1B, V1C, and V0d subunits (Figure S1). These tagged subunits were individually ectopically expressed in amoebic trophozoites under the control of a tetracycline-inducible system. Stable expressing trophozoites were generated and the expression and subcellular localization of each subunit were determined using immunoblotting and immunofluorescence confocal microscopy, respectively (Figure 1A, B, C). We observed that all the tested V-ATPase subunits were expressed upon tetracycline induction. Our subcellular localization studies showed that V-ATPase subunits are highly enriched at the cell surface. In addition, we also observed that these subunits are localized to intracellular vacuolar compartments. However, fluorescent intensities on these objects were low when compared with cell surface. Based on these observations, we hypothesized whether *Entamoeba* V-ATPase localizes to the plasma membrane, as reported in other organisms [40, 41]. The V0a subunit of V-ATPase is the largest membrane-embedded component, essential for core proton translocation across cellular membranes [42]. Therefore, we generated a polyclonal antibody against a synthetic peptide corresponding to the predicted extracellular region of *Entamoeba* V0a subunit. Our detailed fixed, live immunofluorescence analysis and antigen competition assay showed antibody specificity to V0a antigen in native conformation (Figure S2). However, a clear band was shown by the V0a antibody only in the membrane-rich fractions. Further, this antibody was utilized in wild-type trophozoites to visualize endogenous localization of V0a subunit along with a known plasma membrane lipid phosphatidylinositol 4,5-bisphosphate (PIP2) [43]. Our confocal images suggest that the V0a subunit is highly enriched at the cell surface and colocalizes with PIP2 (Figure 1D). Further, HA-V1A subunit-expressing trophozoites were probed with anti-PIP2 antibody and identified that the V-ATPase subunit V1A and PIP2 also colocalizes at the cell surface (Figure 1E). Further, to support the presence of V-ATPase at the plasma membrane, a cell surface biotinylation approach was used to separate the plasma membrane pool from the trophozoite [44, 45]. The HA-V1A catalytic subunit-expressing trophozoites were biotinylated and the isolated plasma membrane fractions were used for immunoblotting. Our results clearly showed that the HA-V1A subunit was detected along with plasma membrane receptor, heavy chain of Gal/GalNAc lectin (Hgl) subunit [46] in the biotinylated fractions (Figure 1F). It is very well known that V-ATPase enriches on the lysosomal compartments to promote acidification and lysosomal function [47]. Therefore, we probed HA-V1A and HA-V0d trophozoites with a widely used lysosome marker, LysoTracker. Previously, it has been shown that LysoTracker labels the acidified endosomes and lysosomes in *E. histolytica* [48, 49]. We found that both HA-V1A and HA- V0d are localized on the LysoTracker positive compartments (Figure S3). Our quantification analysis revealed that HA-V1A trophozoites contained approximately 53.9% more acidic vacuoles compared to HA-V0d trophozoites. Further, we calculated the association between V-ATPase subunits and LysoTracker. Our results indicate that the HA-V1A vacuoles showed more association with LysoTracker (36.1 ± 14.7%, n= 722 LysoTracker vesicles/20 trophozoites) when compared with HA-V0d associated vacuoles (13.0 ± 6.2%, n= 261 LysoTracker vesicles/20 trophozoites). Together, these results suggest that the HA-tagged V-ATPase subunits are expressing in amoebic trophozoites and they are associated with cell membrane as well as intracellular acidic compartments.

**Figure 1.**
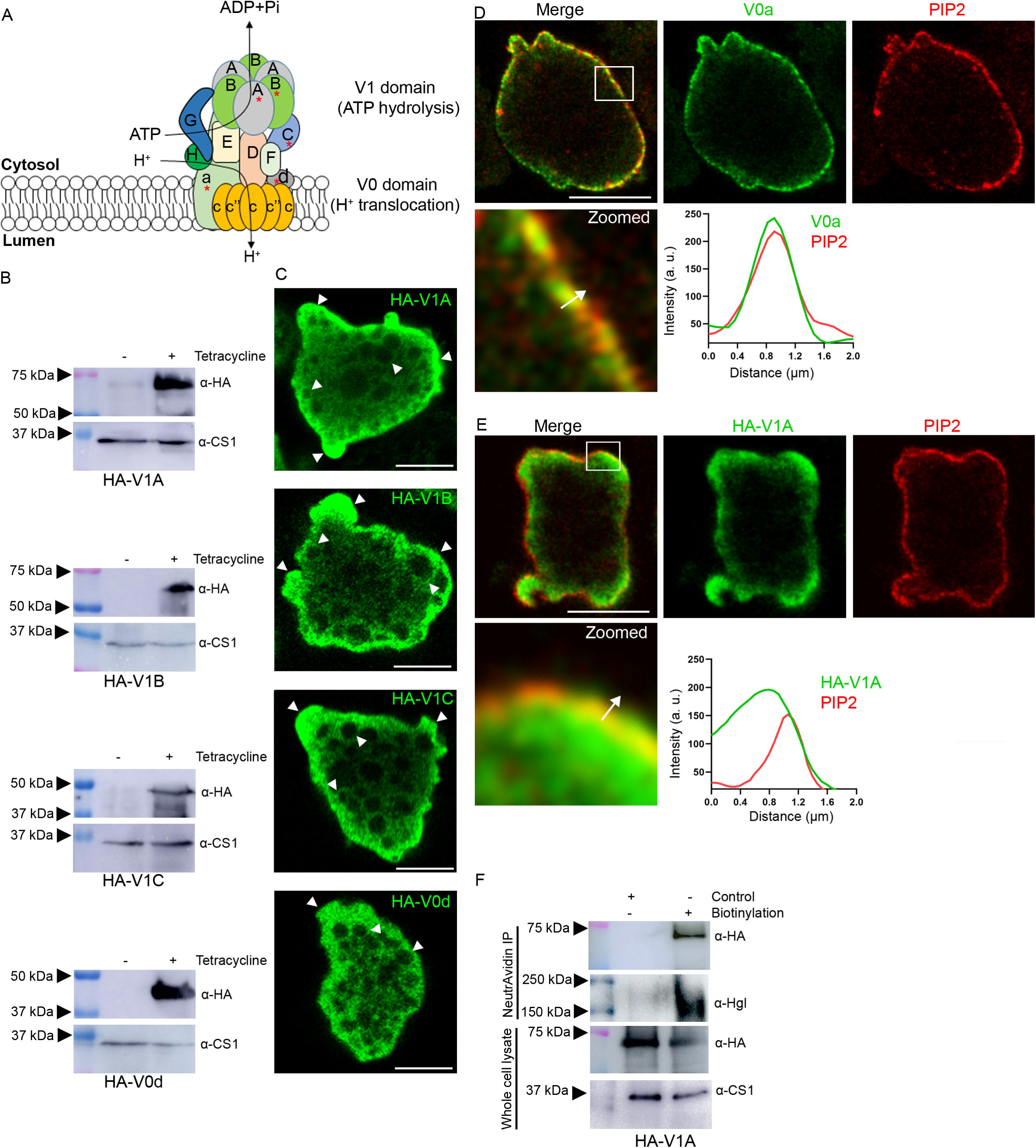
Expression and cell surface localization of *E. histolytica* V-ATPase subunits. **A.** An outline of the V-ATPase complex, *red* asterisk highlights the subunits analyzed in this study (HA-V1A, HA-V1B, HA-V1C, HA-V0d and endogenous V0a). **B.** Trophozoites stably expressing V-ATPase subunits (HA-V1A, HA-V1B, HA-V1C, and HA- V0d) were grown in absence and presence of tetracycline and lysed; 40 µg of whole cell lysate was subjected to immunoblotting with anti-HA and anti-cysteine synthase 1 (CS1) antibodies. **C.** Tetracycline induced trophozoites expressing individual HA-tagged V-ATPase subunits (HA-V1A, HA-V1B, HA-V1C, and HA-V0d) were incubated on glass multi-well slides for 15 minutes at 37°C. Following incubation, trophozoites were fixed, permeabilized and processed for immunofluorescence assay using an anti-HA antibody. *White* arrowheads indicate the distribution of V-ATPase subunits. Scale bars, 10 µm **D.** Wild-type amoebic trophozoites (untransfected) were incubated on glass multi-well slides for 15 minutes at 37°C. The trophozoites were fixed and processed for immunofluorescence assay using *Entamoeba* anti-V0a and anti-PIP2 antibodies. The *white* square object represents a zoomed panel. The line intensity plot indicates the colocalization between V- ATPase V0a subunit and PIP2 across the *white* arrow at cell surface. Scale bars, 10 µm **E.** HA-V1A expressing trophozoites were incubated for 15 minutes on glass surface and probed with anti-HA and anti-PIP2 antibodies. The *white* square object represented into a zoomed panel. The line intensity plot indicates the colocalization between V-ATPase V1A subunit and PIP2 across the *white* arrow at cell surface. Scale bars, 10 µm **F.** The cell surface proteins of HA-V1A trophozoites were biotinylated and immunoprecipitated using NeutrAvidin beads. The elutes were resolved by SDS-PAGE, immunoblotted using anti-HA and anti-Hgl antibodies. The Hgl was used as a positive control and CS1 was used as a loading control in the NeutrAvidin bound fraction and whole cell lysate, respectively.

### During macropinocytic cup formation, V-ATPase complex transiently recruits and subsequently dissociates from nascent macropinosome

Previously, it has been shown that V-ATPase controls the transport of cholesterol to the plasma membrane that regulates macropinocytosis in mammalian cells [37]. Our imaging data suggests that V-ATPase subunits localize to pseudopod-like structures that primarily consist of active membrane trafficking zone. Therefore, we sought to ask whether V-ATPase is associated with the F-actin network at the early stage of macropinocytosis. The V-ATPase HA-V1A subunit expressing trophozoites, under steady-state conditions were probed with anti-HA and phalloidin (F-actin). Surprisingly, we observed that the V1A subunit co-localizes with F-actin at macropinocytic cups (Figure 2A). Similar observations were also made for other V1 domain subunits (HA-V1B and HA-V1C). Further we wanted to confirm whether the holo V-ATPase complex recruits at macropinocytic cup. Membrane-associated V0d subunit was known to be implicated in the V-ATPase assembly and scaffolding of the V1 and V0 domains [27]. The HA-V0d subunit expressing trophozoites were probed with anti-HA and phalloidin. Our confocal images clearly indicated that the V0d subunit indeed localizes at the macropinocytic cup (Figure 2A). Further to see spatial distribution of V-ATPase, HA-V1B expressing trophozoite images were 3D reconstructed. Our image analysis revealed that HA-V1B was enriched predominantly at the rim of the cup (Figure 2B). Since, the tested V-ATPase subunits were colocalized with F-actin at pinocytic cups, we further analyzed their colocalization throughout the trophozoites using Aivia image analysis software (Figure S4). Interestingly, we found that V1B showed highest levels of Pearson’s correlation coefficient (*r*) (0.41 ± 0.04, n = 600 trophozoites) followed by V1A (0.31 ± 0.07, n= 600 trophozoites), V1C (0.29 ± 0.12, n = 600 trophozoites) and V0d (0.23 ± 0.07, n = 600 trophozoites). Next, we asked whether the overexpression of V-ATPase subunits affects the biogenesis of macropinocytic cups. To address this, HA-V1A, HA-V1B, HA-V1C, and HA-V0d expressing trophozoites were incubated for 15 minutes under steady-state conditions, and macropinocytic cups were quantified. Interestingly, our results showed that V1A and V1B overexpression significantly upregulated the macropinocytic cup formation by 76.80% and 43.02% respectively, when compared with the vector control (Figure 2C). In contrast, V1C and V0d expressing subunits led to a slight increase in the cup numbers, but the difference was statistically insignificant relative to the vector control. Together, these results indicate that the V-ATPase is recruited at macropinocytic cups and the overexpression of catalytic subunits elevates the overall cups formation.

**Figure 2.**
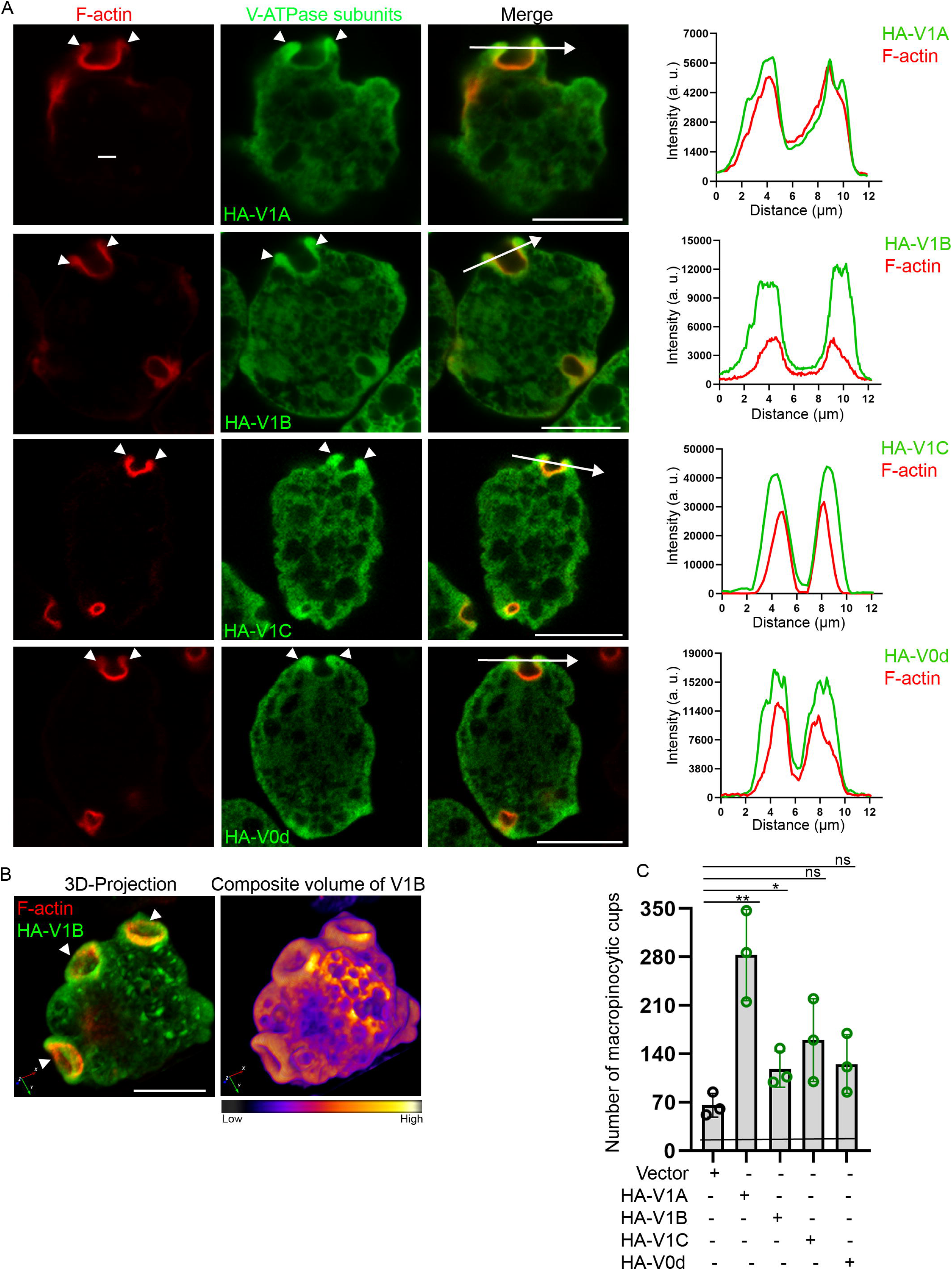
V-ATPase subunits localizes to the macropinocytic cups. **A.** Trophozoites expressing individual HA-tagged V-ATPase subunits (HA-V1A, HA-V1B, HA-V1C, and HA-V0d) were incubated on glass multi-well slides for 15 minutes at 37°C. Following incubation, trophozoites were processed for immunofluorescence using an anti-HA antibody and phalloidin (F-actin). *White* arrowheads indicate enrichment of V-ATPase subunits at the edge of macropinocytic cup. The line intensity plot indicates the colocalization of V-ATPase subunits and F-actin across the *white* arrow at macropinocytic cup. Scale bars, 10 µm **B.** 3D-projection of macropinocytic cups. 3D-respresentation of the trophozoite displayed multiple macropinocytic cups. The *green* channel shows HA-V1B and *red* channel indicates F-actin. *White* arrowheads show the cups (left side). The same trophozoite was further processed for volume rendering to highlight *green* channel intensity at the cups (right side). Scale bar, 10 µm **C.** Quantification of macropinocytic cups. Amoebic trophozoites stably expressing the V- ATPase subunits (HA-V1A, HA-V1B, HA-V1C, and HA-V0d), were incubated for 15 min at 37°C under steady state condition. Post incubation, trophozoites were processed for immunofluorescence using phalloidin. The *Z*-stack of confocal images were acquired and number of macropinocytic cups were counted manually. Bar graph represents the mean ± standard deviation (SD) of three independent experiments (500 trophozoites/each experiment). Statistical significance was determined by unpaired Student’s t-test (\*\**p*<0.01, 0.0053; \**p*<0.05, 0.0444; ns: non-significant)

Further, we asked whether the V-ATPase complex is dynamically recruited at the macropinocytic cup and assembled at the site of cup initiation. First, we employed HA-V1A subunit-expressing trophozoites and probed them with endogenous anti-V0a and F-actin, and imaged the macropinocytic cups. Our results clearly showed the enrichment of both V1 and V0 subunits at the cup, providing another layer of evidence for the V-ATPase complex recruitment in this process (Figure 3A). To further investigate the spatiotemporal recruitment of V-ATPase during macropinocytic cup formation, we generated GFP-V0d expressing trophozoites and confirmed with western blotting (Figure 3B). Subsequently, GFP-V0d expressing trophozoites were subjected to dextran uptake assay and the dynamics were visualized. Our live-cell imaging data of the GFP-V0d trophozoites clearly showed that V-ATPase enrichment occurs prior to the formation of negative curvature and increases along the membrane ruffles. Later, the V0d enriched ruffles seal and trap the dextran within the vacuoles called macropinosomes (Figure 3C and Supplementary Movie 1). The fluorescence intensity suggests that V0d is peaked at the moment of cup formation, then rapidly declines upon the closure of cup (18.99 seconds) and further reduces on the nascent macropinosome (20-21 seconds) (Figure 3C and Supplementary Movie 1). Next we quantified the GFP-V0d fluorescence intensities on cups and pinosomes in live trophozoites as depicted in the cartoon (Figure 3D left side). Our quantification data showed that fluorescence intensities were significantly reduced by 34.07% on macropinosomes when compared with pinocytic cups (Figure 3D right side). Further, we asked whether the V-ATPase complex continues to remain after the closure of pinocytic cup. Previously, it has been shown that F-actin rings (3- 5 µm) form macropinosomes in this parasite [50]. These structures are temporal and actin dissociation occurs within minutes after gulping the extracellular fluid [20]. The V0d subunit acts as a scaffold that helps connect the membrane-embedded V0 sector with the cytosolic V1 sector to stabilize the assembly of V-ATPase complex [26]. Therefore, we investigated the abundance of V0d on macropinocytic cups and actin-positive macropinosomes in the same trophozoite. The HA-V0d expressing parasites were probed with F-actin and were imaged. We selectively monitored the trophozoite that displayed both macropinocytic cups and macropinosomes in different *Z*-planes to avoid discrepancies in expression levels among different trophozoites. The line intensity plots suggest that the V-ATPase V0d abundance is low on the macropinosome (*Z*-plane 5) compared with macropinocytic cup (*Z*- plane 12) in the same trophozoite (Figure 3E). Collectively, these results suggest that V-ATPase complex assembly occurs at macropinocytic cups and disassembles on macropinosomes.

**Figure 3.**
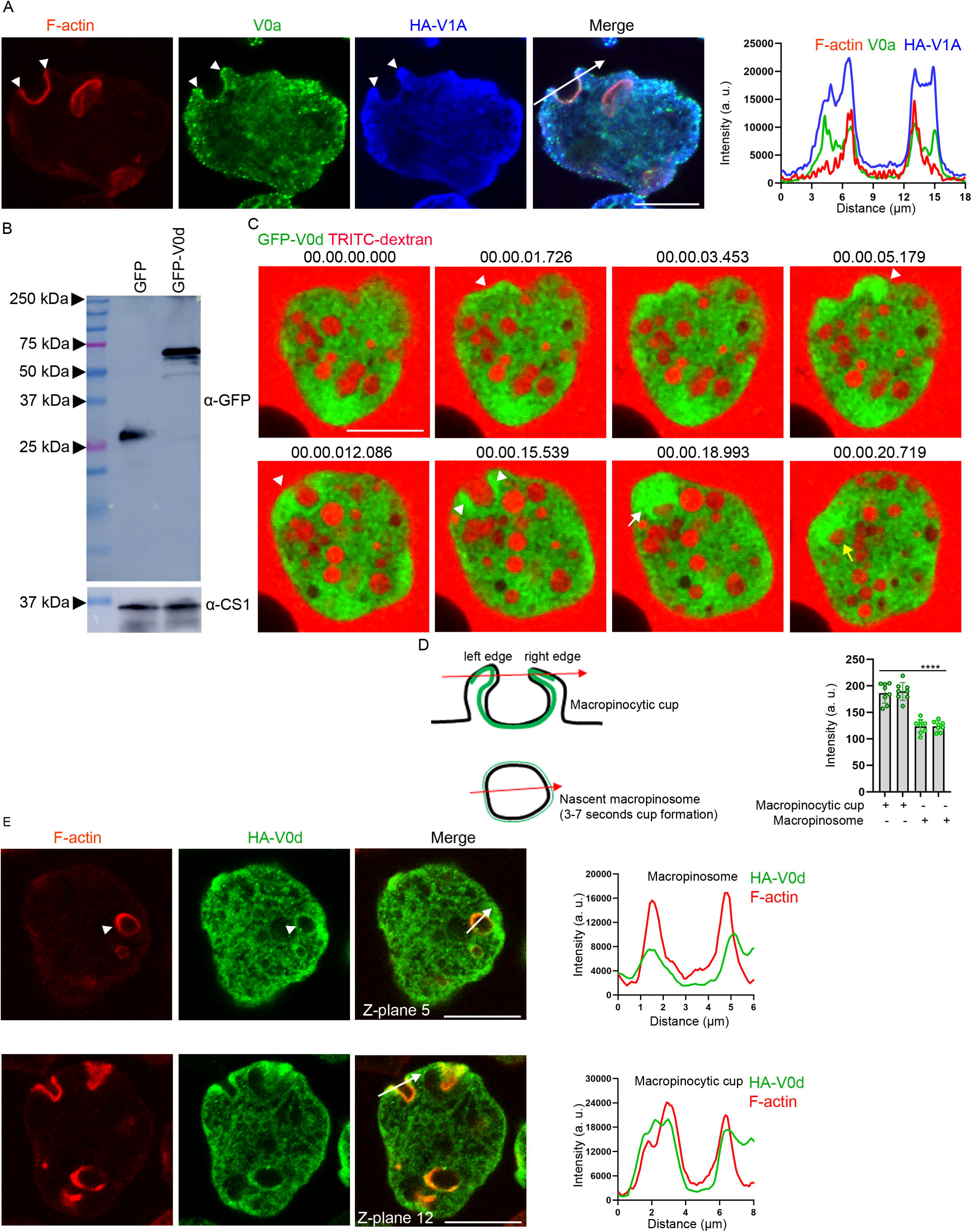
V-ATPase assembles at macropinocytic cups and dissociates from nascent macropinosomes. **A.** HA-V1A expressing trophozoites were incubated for 15 minutes on glass slide and probed with anti-HA, *Entamoeba* anti-V0a and phalloidin. The arrowheads indicate the colocalization of V-ATPase subunits at the pinocytic cups. The line intensity plot indicates V-ATPase subunits and F-actin intensities across *the white* arrow at macropinocytic cup. Scale bar, 10 µm **B.** GFP (vector control) and GFP-V0d expressing trophozoites were lysed and 40 µg of whole cell lysate was subjected to immunoblotting with anti-GFP and anti-CS1 antibodies. **C.** Amoebic trophozoites, transfected with GFP-tagged V-ATPase subunit V0d (*green*), were incubated with 2 mg/ml TRITC-dextran (*red*). Representative frames from time-lapse confocal imaging of GFP-V0d highlight dynamics of macropinocytosis. *White* arrowheads indicate macropinocytic cup invaginations, while the *white* arrow marks closing of the cup. The *yellow* arrow shows reduced abundance of GFP-V0d on the macropinosome. Scale bar, 10 µm **D.** GFP-V0d fluorescence quantification at macropinocytic cups and pinosomes. Cartoon depicts the quantification of GFP-V0d at the edges of macropinocytic cups and nascent macropinosomes. Bar graph represent the mean ± standard deviation (SD) of fluorescence intensities for GFP-V0d across the edges macropinocytic cups and macropinosome. Total 8 different cellular events were quantified. Statistical significance was determined by unpaired Student’s t-test (\*\*\*\**p*<0.0001, 0.0001) **E.** HA-V0d expressing trophozoites were incubated for 15 minutes on glass slide and probed with anti-HA and phalloidin. The line intensity plot indicates V-ATPase subunits and F-actin intensities across the *white arrow* at macropinosome (*Z*-plane 5) and macropinocytic cup (*Z*- plane 12). The arrowheads indicate the colocalization of V0d with actin on macropinosome and macropinocytic cup. Scale bars, 10 µm.

### V-ATPase inhibition reduces the macropinocytic cup formation and rate of macropinocytosis

Our results showed that the V-ATPase complex operates at macropinocytic cups. Next we asked whether the V-ATPase complex function is required for actin polymerization on the cups. Therefore, we employed well-established pharmacological inhibitors bafilomycin and concanamycin that selectively inhibit the V-ATPase function by blocking the proton transport [51, 52]. The working concentrations of these inhibitors were already optimized and successfully used with *E. histolytica* in multiple studies [21, 22]. We treated the wild-type trophozoites with 50 nM bafilomycin or 50 nM concanamycin and macropinocytic cups were studied. In *Entamoeba* and *Dictyostelium*, macropinocytic cups typically form a U-shape that are not dictated by particle/cargo [50, 53], unlike the phagocytic cups morphology which is defined by surface property of the particle. However, other shaped cups such as V-shape and W-shape were also observed during macropinocytosis [54]. Collectively, our quantification data (U, V and W-shaped) showed that the V-ATPase inhibition led to a significantly reduced number of macropinocytic cups, which is 43.4% and 39.5% for trophozoites treated with bafilomycin and concanamycin respectively, when compared with untreated trophozoites (Figure 4A, B). Next, we performed scanning electron microscopy to examine the surface topology of pinocytic cups in untreated trophozoites and those treated with bafilomycin or concanamycin. Our imaging results showed that the parasites treated with V-ATPase inhibitors exhibited irregular and poorly defined macropinocytic structures. In contrast, untreated trophozoites displayed well-defined macropinocytic cups with distinct circular ruffles, indicating that inhibitor treatment alters cup morphology compared to controls (Figure 4C). Finally, we asked whether the rate of macropinocytosis in these treated trophozoites is affected. V-ATPase inhibitor treated trophozoites were subjected to dextran uptake for 5, 10 and 15 minutes, respectively. Our fixed cell image quantification data showed that the V-ATPase inhibition markedly reduced the levels of intracellular dextran in amoebic trophozoites with respect to increasing time (Figure 4D, E). Collectively, these findings suggest that inhibition of V-ATPase activity suppresses the biogenesis, alter the topology of macropinocytic cups, and reduce the rate of macropinocytosis.

**Figure 4.**
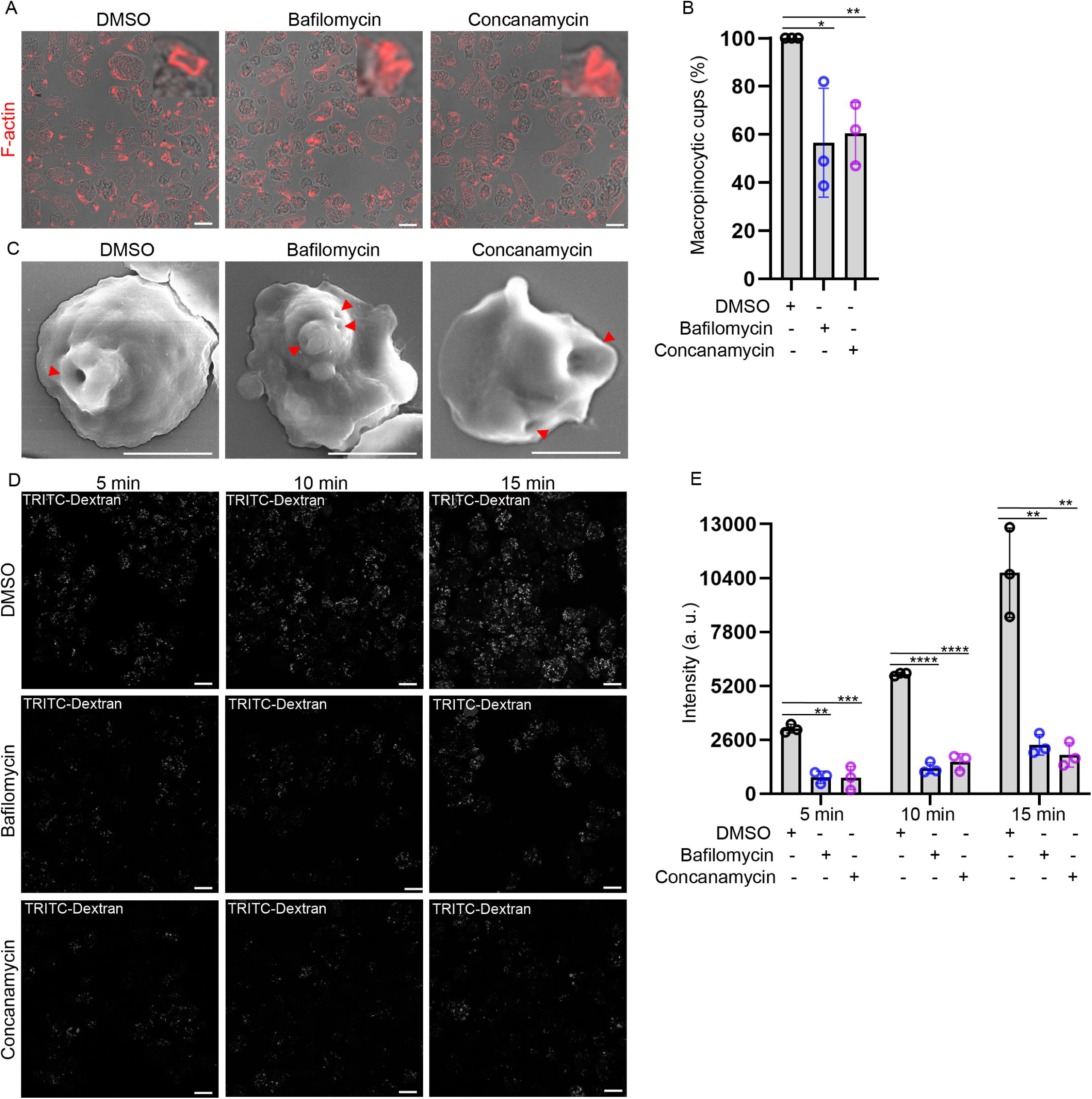
V-ATPase pharmacological inhibition leads to altered morphology of the macropinocytic cups and rate of macropinocytosis. **A.** Wild type amoebic trophozoites were treated with bafilomycin (50 nM) and concanamycin (50 nM) for 1 hour in BIS-33 media. Further, trophozoites were harvested and incubated on glass slide for 15 min at 37°C in BIS-33 media followed by immunofluorescence assay. F-actin was visualised by phalloidin. The inset shows abundant cup-shaped morphologies observed in each condition. Scale bars, 20 µm **B.** Graph represents the number of macropinocytic cups formed in bafilomycin, concanamycin treated and untreated *Entamoeba* trophozoites. Bar graph indicates mean ± standard deviation (SD) of three independent experiments (400 trophozoites/each experiment). Statistical significance was determined by unpaired Student’s t-test (\*\**p*<0.01, 0.0059; \**p*<0.05, 0.0292) **C.** Surface topology of pinocytic cups in amoebic trophozoites treated with bafilomycin and concanamycin were visualized by scanning electron microscopy. Wild type amoebic trophozoites were treated with bafilomycin (50 nM) and concanamycin (50 nM) for 1 hour in BIS-33 media. The *red* arrowheads show cups. Scale bars, 10 µm **D.** Amoebic trophozoites were treated with bafilomycin (50 nM) and concanamycin (50 nM) for 1 hr in BIS-33 media. Further, trophozoites were harvested, incubated with TRITC-dextran (2mg/ml) on glass slide for 5 min, 10 min and 15 min. Later, trophozoites were fixed and imaged using confocal microscope. Scale bars, 20 µm **E.** The fluorescence intensities from TRITC-dextran was measured by using Fiji ImageJ software. Bar graph indicate mean ± standard deviation (SD) of three independent experiments (500 trophozoites/each experiment). Statistical significance was determined by unpaired Student’s t-test for 5 min (concanamycin, \*\*\**p*<0.001, 0.0003; bafilomycin, \*\**p*<0.01, 0.0023), 10 min (\*\*\*\**p*<0.0001, 0.0001) and 15 min (concanamycin, \*\**p*<0.01, 0.0029; bafilomycin \*\**p*<0.01, 0.0024)

### V-ATPase V1B subunit facilitates actin polymerization *in vitro* and its knockdown reduced macropinocytic cup formation

As evident from our quantification (Figure 2C), HA V1A and HA V1B overexpression elevates macropinocytic cup formation. These findings prompted us to investigate whether V1A and V1B subunits are directly involved in macropinocytic cup remodelling through the actin regulatory machinery. We focused on the V1B subunit because it is a highly evolutionarily conserved subunit across eukaryotic kingdoms (Figure 5A) and has been shown to bind with actin [32]. Therefore, GST-V1B recombinant protein was expressed in *E. coli* and purified using affinity-based purification (Figure 5B). First, we wanted to re-establish whether the V1B subunit binds with actin as it is observed with *A. thaliana* V1B subunits. We performed actin co-sedimentation assay to check for actin binding using validated controls. We prepared the F-actin pool and incubated it with α-actinin (positive control), BSA (negative control) or V1B. Subsequently, high speed centrifugation was performed and both soluble and pellet fractions were subjected to SDS-PAGE. Our results showed that V1B co-sediments along with F-actin pellets (Figure 5C, D). These results indicate that V1B binds with F-actin. Interestingly, we also observed enriched G-actin pool along with V1B in the supernatant fractions suggesting either an interaction with G-actin or potentially a severing activity. Next, we asked whether *Entamoeba* V1B polymerizes the actin. However, the role of V1B in direct actin polymerization has not been studied in eukaryotes. We used the G-actin pool and incubated it with α-actinin, BSA or V1B. Remarkably, our results showed that V1B is found in pellet fraction along with polymerized F-actin, indicating that it promotes the polymerization of G-actin to F-actin. (Figure 5E, F). To further analyze the effect of V1B on the kinetics of muscle actin polymerization we performed pyrene-labelled actin polymerization assay. The different concentrations of GST-V1B protein (0.625 - 5.0 µM) were utilized in this assay. Our results demonstrated that V1B enhanced the pyrene fluorescence, suggesting increased actin polymerization. Interestingly, we found that at low concentration V1B (0.625 µM) showed a rapid increase in the fluorescence. However, with increasing concentrations of V1B (1.25 - 5.0 µM) the rate of actin polymerization is declined (Figure 5G), suggesting concentration dependent role in actin dynamics. These results also corroborated with our data (Figure 5C) where V1B showed actin severing activity when it incubated with pool of F-actin. Together, these results suggest that V1B binds with actin and directly polymerize actin *in vitro*. Our previous results showed that pharmacological inhibition of V-ATPase dramatically reduced macropinocytic cups and V1B showed polymerization activity. Therefore, next we sought to ask whether V1B knockdown (KD) impairs macropinocytic cups formation in amoebic trophozoites. The antibody against human V1B was validated against *Entamoeba* lysates, recognizing a predicted ∼55 kDa band (Figure S5). Further, V1B knockdown trophozoites were generated and downregulation was confirmed using western blotting (Figure S6). Under steady state conditions, vector control and V1B depleted trophozoites were probed for F actin. Our image quantification data showed that V1B downregulation significantly reduced the number of macropinocytic cups compared with vector control parasites (Figure 5H). Together, these findings suggest that the V ATPase V1B subunit promotes actin polymerization and is required for macropinocytic cup formation.

**Figure 5.**
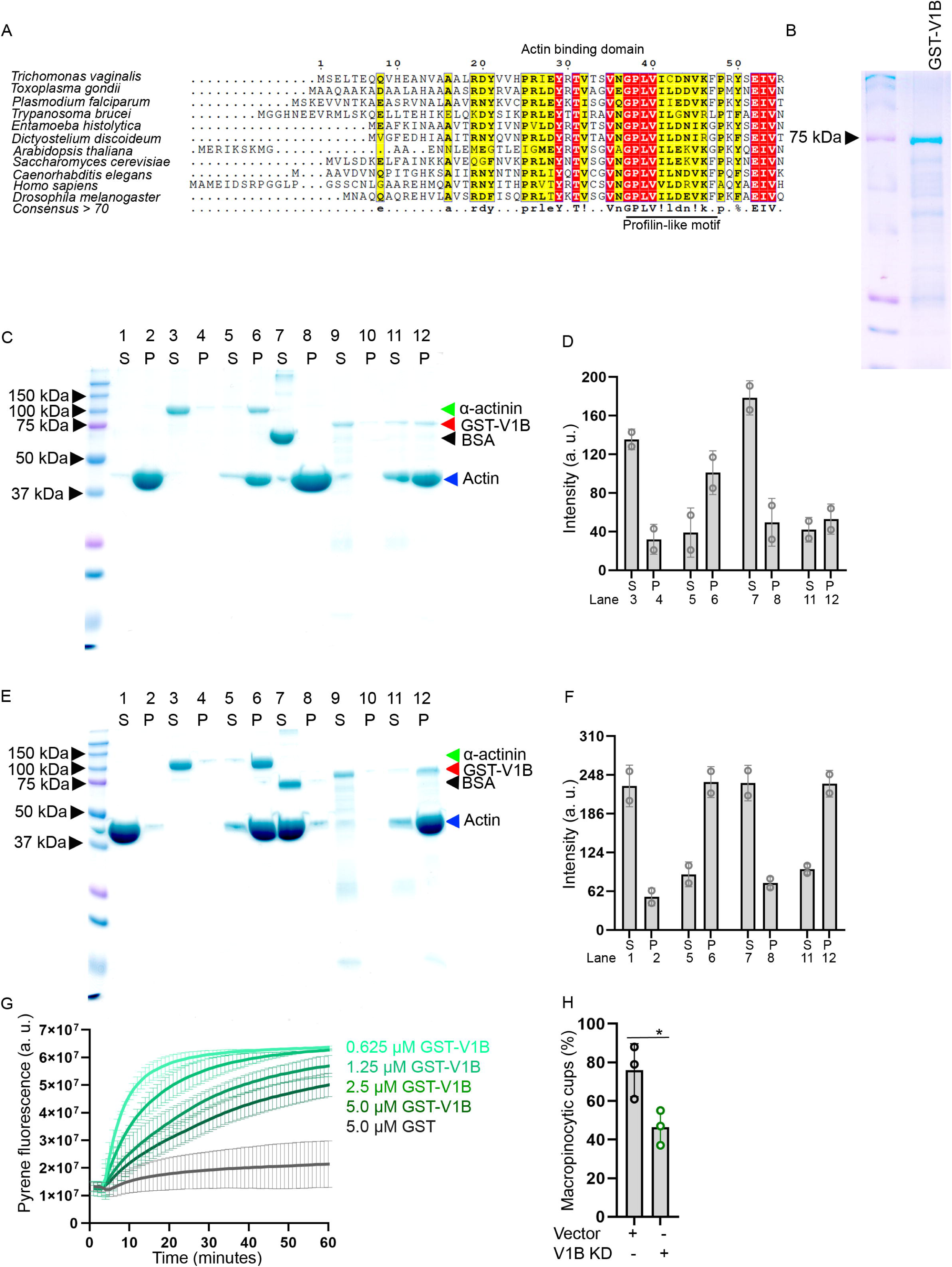
Evolutionarily conserved V-ATPase V1B subunit from *Entamoeba* regulates actin dynamics. **A.** A representation of the multiple sequence alignment of conserved residues within N-terminal actin binding domain with profilin-like motif of the *E. histolytica* V-ATPase, V1B subunit with other eukaryotic species. **B.** Coomassie stained gel image of recombinant GST-V1B (3 µg) purified using glutathione sepharose-based affinity purification. **C.** V1B subunit binds with F-actin. Purified GST-V1B (15 µM), α-actinin (2 µM), BSA (2 µM) were incubated with F-actin for 1 hour and subsequently the reactions were subjected to high speed ultracentrifugation. Supernatant (S) and pellet (P) fractions were collected and analyzed using SDS-PAGE. Arrowheads indicate the bands of GST-V1B (*red*), α-actinin (*green*), BSA (*black*), and actin (*blue*). **D.** The band intensities of α-actinin (lane 3, 4, 5 and 6), BSA (lane 7 and 8) and GST-V1B (lane 11 and 12) were quantified (as labelled in bar graph) using ImageJ. More than 3 independent experiments were performed. The bar graph represents the mean ± standard deviation (SD) from two independent experiments. **E.** V1B subunit polymerizes G-actin. Purified GST-V1B (15 µM), α-actinin (2 µM), BSA (2 µM) were incubated with G-actin for 1 hour and subsequently reactions were subjected to high speed ultracentrifugation. Supernatant (S) and pellet (P) fractions were collected and analyzed using SDS-PAGE. Arrowheads indicate the bands of GST-V1B (*red*), α-actinin (*green*), BSA (*black*), and actin (*blue*). **F.** The band intensities of actin was quantified (as labelled in bar graph) using ImageJ. More than 3 independent experiments were performed. The bar graph represents the mean ± standard deviation (SD) from two independent experiments. **G.** Fluorescence based actin polymerization kinetics was measured using pyrene labelled G-actin. Increasing concentrations of GST-V1B (0.625, 1.25, 2.5 and 5 µM, test) and GST alone (5 µM, negative control) was incubated with pyrene-labelled G-actin and fluorescence was measured at interval of 1 min for period of 60 minutes. The fluorescence intensities were recorded as A.U (arbitrary units) and plotted over time. More than 3 independent experiments were performed. The graph represents the mean ± standard deviation (SD) acquired from the data analyzed from two independent experiments. **H.** Vector control (pTrigger) and V1B knockdown trophozoites were incubated for 15 min at 37°C under steady state condition. Post incubation, trophozoites were processed for immunofluorescence using phalloidin. The *Z*-stack of confocal images were acquired and number of macropinocytic cups were counted. Bar graph represents the mean ± standard deviation (SD) of three independent experiments (500 trophozoites). Statistical significance was determined by unpaired Student’s t-test (\**p*<0.05, 0.0353)

### Phosphatidic acid pulldown reveals the presence of V-ATPase subunits with V1B showing affinity *in vitro* and their interaction modulate actin polymerization

Based on our time-lapse imaging, V-ATPase accumulates prior to the formation of negative curvature of the plasma membrane (Figure 3C). In addition, V1B binds and polymerizes the actin to form macropinocytic cups along with cytoskeleton-lipid regulatory signaling. In mammalian phagocytes, PA plays a crucial role in membrane ruffling and macropinocytosis [2]. In addition, PA acts as pH biosensor [55], regulates the actin polymerization [56] cytoskeleton and signaling pathways, including PI3K, in generating negative curvature of the membrane, and phospholipase D production [57–59]. However, the link between PA and V-ATPase remains understudied in eukaryotes. To explore this, PA-interacting proteins were pulled down from *E. histolytica* trophozoite lysates using PA-conjugated agarose beads and subsequently identified by mass spectrometry (Figure 6A). The unlabelled agarose beads were used as a control. The unique proteins identified in this study can be found in Supplementary Table S3. Our results identified 291 unique proteins which were listed in the PA interactome, representing various classes (Figure 6B and Supplementary Table S4). Interestingly, several V-ATPase subunits (V1A, V1B, V1D and V1E2) were reproducibly found in the PA-pulldown experiments (Figure 6C). In addition, we found several actin-binding, nucleation, polymerization and membrane transport proteins in analysis (Supplementary Table 3). Importantly, our results demonstrated that V1B subunits directly modulate actin polymerization (Figure 5E, F), highlighting a mechanistic link between PA-V-ATPase interactions and cytoskeletal regulation. To further test this interaction, lysates from HA-V1B expressing trophozoites were incubated with PA beads, and bound proteins were analyzed by western blotting. Our results indeed revealed that HA-V1B was pulled down with phosphatidic acid (Figure 7A). To verify whether this interaction is specific to phosphatidic acid *in vitro*, GST-V1B protein was purified and used for cellular lipid assays based on lipid strips immobilized with 100 pmol of various biologically active lipids. Our findings showed that the V1B subunit specifically binds to certain phospholipids, including several phosphatidylinositol mono-/bis-/tri-phosphates, phosphatidylserine and phosphatidic acid (Figure 7B, C). Notably, V1B interactions were reproducible and consistently highest with PA in multiple experiments.

**Figure 6.**
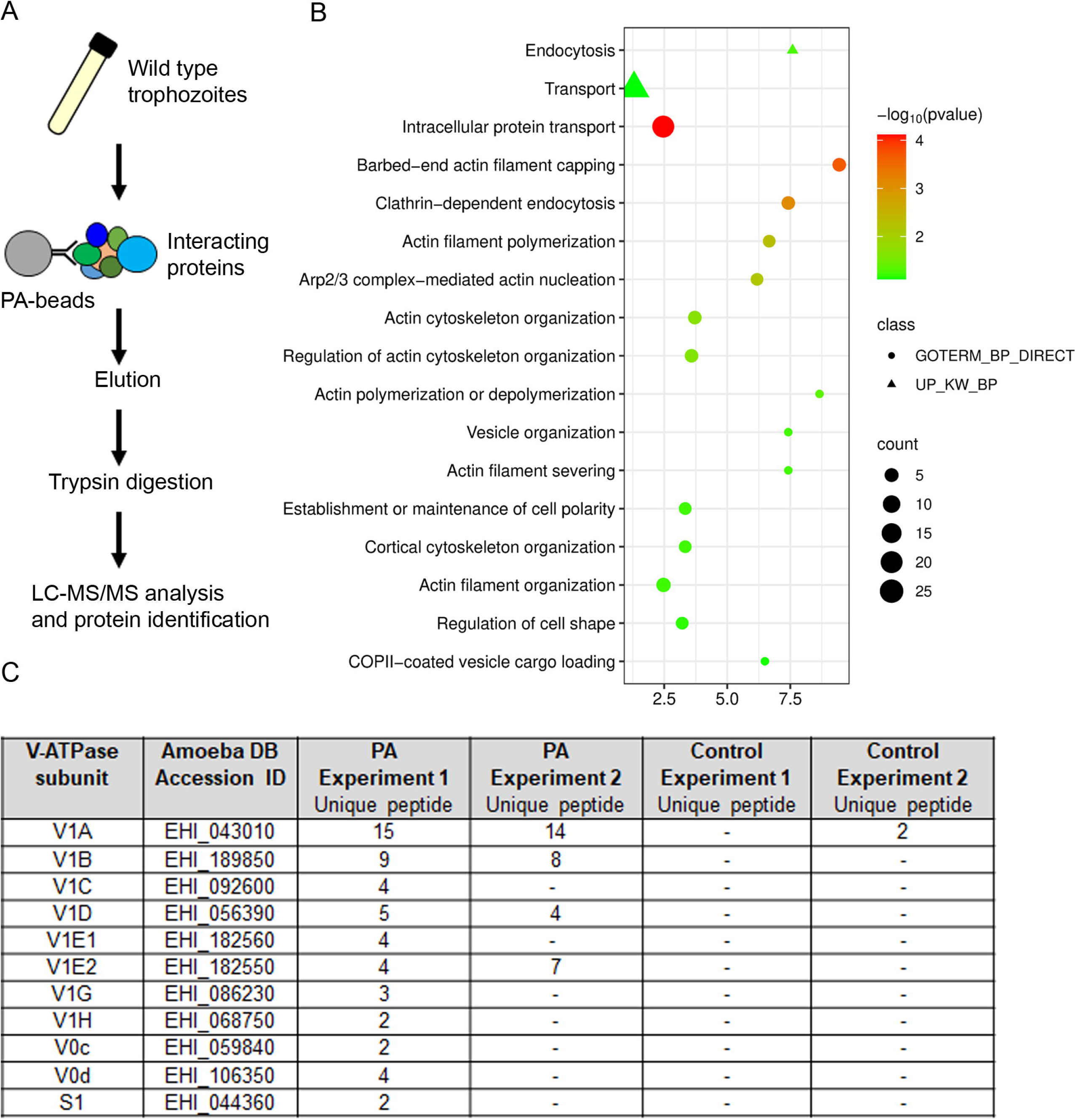
PA pulldown shows multiple V-ATPase subunits. **A.** The depicted workflow is used in identification of PA-associated proteins. *E. histolytica* wild type trophozoites were subjected to pulldown assays with PA-conjugated agarose beads and control beads. The pulldown samples were processed and subjected to mass spectrometry. **B.** Gene ontology enrichment of unique proteins enriched in PA pulldown were represented as an enrichment bubble blot with bubble color corresponding to the statistical significance – log_10_ (*p*-value) and bubble size representing the number of proteins per biological process. **C.** The table represents the peptide count of V-ATPase subunits identified under the proton transmembrane transport in two independent experiments.

**Figure 7.**
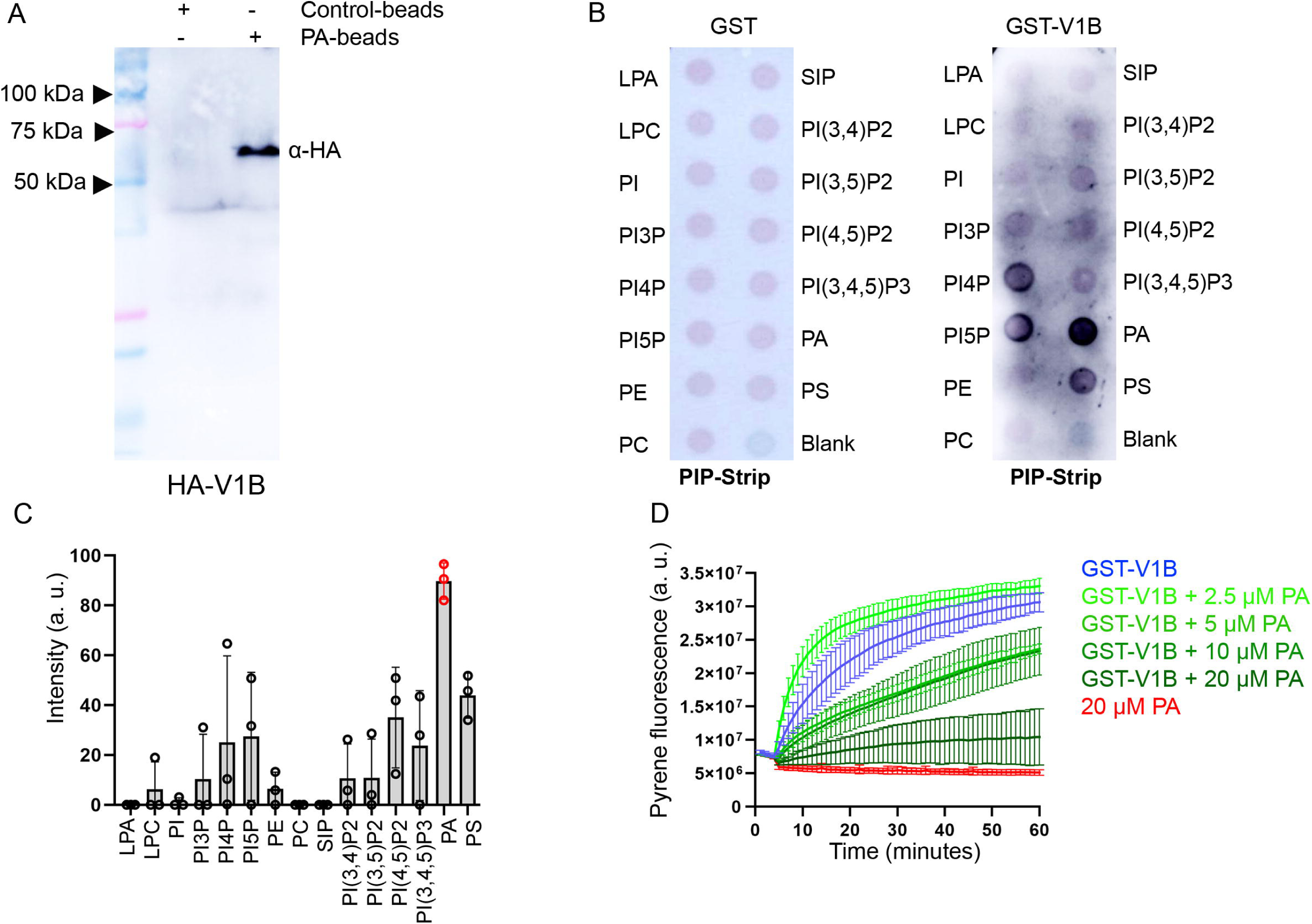
V1B reproducible binds with PA and their interaction modulate actin polymerization. **A.** PA-conjugated beads pulldown the V1B. HA-V1B expressing trophozoites lysate were incubated with PA-beads or control beads. The bound proteins were subjected to western blotting and probed with anti-HA. B. Lipid binding specificity of V1B. The 7 µg/ml of GST alone (as control left side) and 7 µg/ml GST-V1B (right side) incubated on the PIP-strips spotted with 100 pmol of each lipid as indicated and probed with anti-GST. PA, phosphatidic acid; PC, phosphatidylcholine; PE, phosphatidylethanolamine; PI, phosphatidylinositol; PI(3)P, phosphoinositide 3 phosphate; PI(4)P, phosphoinositide 4 phosphate; PI(5)P, phosphoinositide 5 phosphate; PI(3,4)P2, phosphoinositide 3,4 bisphosphate; PI(3,5)P2, phosphoinositide 3,5 bisphosphate; PI(4,5)P2, phosphoinositide 4,5 bisphosphate; PI(3,4,5)P3, phosphoinositide 3,4,5- triphosphate; PS, phosphatidylserine; LPA, lysophosphatidic acid; LPC, lysophosphocholine; S1-P, sphingosine 1-phosphate. C. The densitometric quantification of lipid interaction with GST-V1B was measured by calculating the relative intensities of each lipid spot using ImageJ software. Bar graph represents the mean ± standard deviation (SD) acquired from the data analyzed from three independent experiments. D. Fluorescence based actin polymerization kinetics was measured using pyrene labelled G- actin. V1B (0.625 µM) alone and in the presence of increasing concentrations of PA (2.5, 5, 10 and 20 µM) was incubated with pyrene-labelled G-actin and fluorescence was measured at interval of 1 min for period of 60 minutes. The fluorescence intensities were recorded as A.U (arbitrary units) and plotted over time. The graph represents the mean ± standard deviation (SD) acquired from the data analyzed from two independent experiments.

Further, we sought to ask whether PA-binding to V1B can directly modulate actin polymerization activity, or these interaction has independent functions. Previously, it has been shown that *Arabidopsis* capping protein binds with PA and module actin polymerization [60]. To test with V1B, different concentrations of PA (2.5, 5, 10 and 20 µM) were used along with V1B (0.625 µM) in pyrene-actin polymerization assays. Our results showed that with increasing concentration of PA (5, 10 and 20 µM) along with V1B resulted in decreased rate of actin polymerization (Figure 7D). Interestingly, we found that lowest concentration of PA (2.5 µM) along with V1B showed higher rate of actin polymerization when compared with V1B alone. Collectively, these results suggest that V1B binds with PA and directly affect the rate of actin polymerization.

## Discussion

This study unveils four major findings. First, a major pool of *Entamoeba* V-ATPase localizes on the cell surface. Second, V-ATPase complex dynamically recruited on macropinocytic cups and dissociates from nascent macropinosome. Third, the V-ATPase subunit V1B uniquely polymerizes actin and regulates macropinocytic cups formation. Lastly, PA, a pH biosensor, pulls down V-ATPase and actin-polymerizing proteins, with V1B driving the actin polymerization.

The canonical function of V-ATPase is to regulate organelle acidification and maintain intracellular pH homeostasis. First, our results showed that the V-ATPase complex localizes to the plasma membrane and intracellular compartments. Previous studies postulated the presence of *Entamoeba* V-ATPase on the plasma membrane [61, 62]. Our results corroborate previous finding where V-ATPase subunits were identified with plasma membrane fractions of *E. histolytica* in mass spectrometry analysis [45]. The localization of V-ATPase in the plasma membrane implies that the parasite uses this proton pump to regulate extracellular acidification, drive motility and invade host tissue [40, 63]. A few seminal studies on axenically cultured and clinically isolated *Entamoeba* trophozoites demonstrated that their surface-active lysosomes can be triggered by host cell responses [64, 65]. Interestingly, the plasma membrane localization of V-ATPase is specific to cell types. The kidney intercalated cells, bone osteoclast cells, epididymis clear cells and various cancer cells uses plasma membrane V-ATPase to acidify the external environment [66–68]. Furthermore, it would be interesting to study whether *Entamoeba* plasma membrane V-ATPase is required for extracellular acidification and how it might contribute to pathogenesis. Our study uncovers a previously unrecognized role for V-ATPase subunits in the early stages of macropinocytosis, expanding the classical view of V-ATPase as a late endosomal acidifier. The dynamic recruitment of V-ATPase to macropinocytic cups and its co- localization with actin suggest that V-ATPase is actively involved in shaping the actin architecture required for cup formation and closure. These findings are consistent with prior reports demonstrating that V1B and V1C subunits can directly bind to F-actin and G-actin thereby influencing filament stability and organization [30, 31]. In *Manduca sexta*, V1C was shown to cross-link actin filaments and enhance polymerization, supporting a structural role in cytoskeletal remodelling [33]. We also observed that V-ATPase assembles upon the formation of pinocytic cup and disassembles from the nascent pinosomes. Phosphatidylinositol 3 kinase (PI3K) and nutrient-sensing activities are important for V-

ATPase assembly in eukaryotes [69, 70]. Previous studies demonstrate that *E. histolytica* uses PI3K signaling for actin polymerization to control the shaping of various templates of endocytosis [20, 71, 72]. It can be hypothesized that PI3K signaling converts PIP2 to PIP3 which drives actin-mediated cup formation [17, 54]. Previously it has been shown that the *Entamoeba* PIP3 lipid biosensor FYVE (Fab1p, YOTB, Vac1p and EEA1) finger protein domain localizes to phagocytic cups, suggesting the contribution of PIP3 signaling in this process [73]. Based on our results, V-ATPase V0d dissociates post 18 seconds after a gulp of dextran. While previous results showed that the actin crown is released from the macropinosome after approximately 35 seconds post internalization [19]. These results suggest that V-ATPase disassembly initiates prior to actin removal from the macropinosome. It can be speculated that, possibly phosphoinositide-rich micro-domains alteration may lead to re-localization of V-ATPase assembly at plasma membrane to maintain a pool of V-ATPase for further rounds of macropinocytosis. Further, it would be interesting to know what are the factors that may influence the disassembly of V-ATPase from macropinosomes. We found that V-ATPase is dynamically recruited to macropinocytic cups and dissociates/disassembles from the macropinosomes. These results imply that macropinosomes halt or pause before further maturation. In the past study, it has been shown that dextran-filled macropinosomes take at least 2 hours to neutralize their contents in *E. histolytica* (Meza 2004). This observation raises the possibility of the reassembly of V-ATPase during the maturation of endosomes. It has been supported that V-ATPase function is critical for phagosome maturation in *Entamoeba* species [22]. An alternate possibility is that rapid acidification may occur at very early stage upon the formation of the macropinocytic cup due to V-ATPase assembly at that site. In another context, it has been observed that disassembly and assembly of V-ATPase is necessary to regenerate lysosomes in continuously fed cells [47].

Our results demonstrated that pharmacological inhibition of V-ATPase markedly reduced the number of macropinocytic cups, rate of macropinocytosis and altered cup surface topology, underscoring its functional importance in this nutrient acquisition pathway. Previously it has been shown that the V-ATPase inhibitor bafilomycin increases the pH of pinosome and reduces the pinocytosed volume in *E. histolytica* [61]. These results suggest that the V-ATPase assembly and disassembly regulation is needed at the site of negative curvature during macropinocytosis. It has been shown that V-ATPase inhibitors block the stable complex conformation without transporting the H^+^ ion [52, 74]. Further, our findings showed that V1B knockdown significantly impaired the pinocytic cup formation. The reduced number of cups is possible in multiple ways. First, V-ATPase V1B subunit contributing directly to actin polymerization. Alternatively, disruption of V1B is likely to influence V-ATPase complex formation and proton translocation, ultimately modulating the local pH at the plasma membrane. In mammalian cells it has been tested that the submembranous pH cannot be controlled by V-ATPase in Ras-induced macropinocytosis [37]. However, it cannot be ruled out whether the global pH alteration leads to this phenotype.

We observed that overexpression of V1A and V1B subunits showed significantly high number of cups among other tested subunits. This result implies that V1A and V1B act as a catalytic sector by which ATP hydrolysis occurs and mechanical energy is employed for rotation of the turret and transport of the H^+^ ions. Nevertheless, we cannot rule out the possibility that these subunits scaffold the actin polymerization and stability during this process. The V1B subunit of the cytosolic V1 domain is a highly conserved component of the V-ATPase pump that not only couples ATP hydrolysis to proton translocation but also interfaces with the actin cytoskeleton in ways that have diversified across eukaryotic evolution. We identified that the V-ATPase V1B subunit is highly conserved among other subunits of V-ATPase (Table S1). Interestingly, our data showed for the first time that V1B polymerizes G-actin to F-actin. However, increasing concentration of V1B slightly slows down the rate of polymerization, suggesting its biphasic role in actin dynamics. Moreover, *Arabidopsis* V1B proteins bind with actin and stabilize the actin filaments but direct polymerization activities were not observed [32]. Taken together, these findings show that the V1B subunit, beyond its ancestral role in actin binding, has evolved the capacity to polymerize actin, highlighting the evolutionary diversification of V-ATPase.

The interaction between V-ATPase subunits and PA adds a compelling layer to this mechanism. PA is known to modulate actin polymerization, membrane curvature and recruit actin-binding proteins [56, 57]. Our unbiased mass spectrometry findings also revealed that PA serves as a lipid scaffold for V-ATPase and an actin polymerization hub. Further, we identified that the V1B subunit polymerizes actin and reproducibly binds with PA. The V1B subunit contains an N-terminal conserved profilin-like motif that has already been shown to bind with actin. In humans, testis-specific profilin isoform 4 binds with PA [76], suggesting that PA-interaction is coupled with actin binding proteins. Notably, V1B actin polymerization activity was positively regulated by lower concentrations of PA, but negatively influenced by higher concentrations. Previously, it has been shown that higher concentrations of PA sequential reduce actin filament assembly function of *Arabidopsis* capping protein [60]. It can be assumed that PA abundance could serve as a regulatory switch during different types of endocytosis. While a small group of eukaryotic PA-binding proteins has been documented [77]. Further investigation is required to map the precise PA-binding sites in *Entamoeba* V1B. PA acts as a pH biosensor and is linked with V-ATPase in yeast metabolism [55]. In yeast, PA biosynthesis via Pah1p phosphatidic acid phosphatase influences V-ATPase disassembly and vacuolar acidification [78]. In mammalian neuroendocrine cells, PA generated by phospholipase D facilitates the interaction of V-ATPase subunit/s with ARF nucleotide-binding-site opener (ARNO) and Arf6, thereby promoting localized acidification and membrane fusion [79]. Based on our result, multiple V-ATPase subunits are pulled down with PA, suggesting a coordinated lipid-protein interface that could regulate both proton pump activity and actin dynamics in a spatially restricted manner. Furthermore, it would be interesting to investigate whether individual V-ATPase subunits show specific affinities with selective lipid species to play distinct roles during various endocytic processes [80].

In this study, we uncover that V-ATPase dynamically regulates macropinocytic cup remodelling via a PA-mediated signaling cascade that drives actin polymerization (Figure 8). This study underscores the need to investigate whether higher eukaryotes preserve this pathway to fuel nutrient acquisition through endocytosis. Furthermore, our work establishes a foundation for repurposing V-ATPase-specific inhibitors as targeted therapeutics against the enteric pathogen *E. histolytica*.

**Figure 8.**
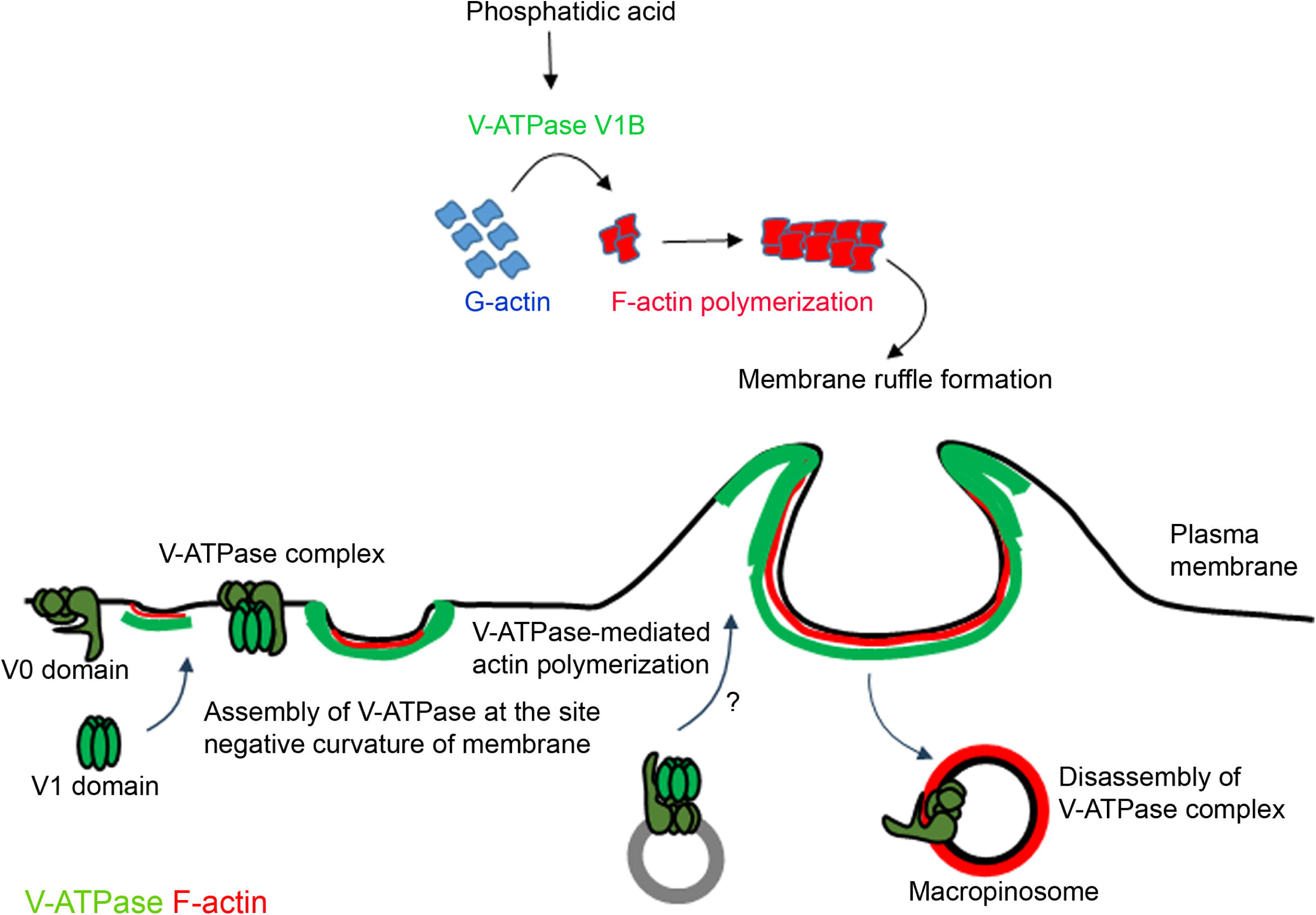
Proposed model for V-ATPase-mediated regulation of macropinocytosis in *E. histolytica*. This model depicts the coordinated role of V-ATPase and phosphatidic acid in regulating actin dynamics during macropinocytosis. V-ATPase rapidly accumulates at the plasma membrane through local assembly of the complex prior to macropinocytic cup formation. The V1B subunit binds and polymerizes the actin along with scaffolding the cytoskeletal proteins. PA functions as a pH-sensitive lipid mediator that binds V1B subunit and facilitates actin polymerization. Upon cup closure, V-ATPase dissociates from the nascent macropinosome.

## Methods

### E. histolytica cell culture

*E. histolytica* HM-1:IMSS trophozoites were grown axenically in BIS-33 medium supplemented with 2% (v/v) Diamond’s vitamin mix, 15% (v/v) heat-inactivated adult bovine serum (RM10913, HiMedia Labs, India), and 1X Antibiotic-Antimycotic (15240062, Gibco, USA). The trophozoite cultures were grown at 35.5°C for routine culturing.

### Cloning of V-ATPase subunits into amoebic expression vectors

Total RNA from the *E. histolytica* wild-type trophozoites was isolated using the Purelink RNA mini kit (12183020, Invitrogen, USA). cDNA was synthesized from the total RNA using the High-Capacity RNA-to-cDNA™ Kit (4387406, Invitrogen, USA) and was further used as a template for amplifying the V-ATPase subunit genes [V1A (EHI_043010), V1B (EHI_189850), V1C (EHI_092600) and V0d (EHI_106350)] with gene specific-primers (Supplementary Table 2). PCR amplification was performed using Phusion High-Fidelity PCR master mix (M0531S, New England Biolabs, USA). The amplified PCR products were resolved on 1% agarose gel by electrophoresis at 90V for 1-2 hours, visualized under UV transilluminator and the bands corresponding to the expected size were excised and purified using the QIAquick Gel Extraction Kit (28704, Qiagen, Germany). The amoebic expression vectors pEhTex-HA and pEhEx-GFP as well as the amplified PCR products (inserts) were double digested with *SmaI* and *XhoI* restriction enzymes (R0141S, R0146S; New England Biolabs, USA) and dephosphorylated with calf intestinal alkaline phosphatase (M0525S, New England Biolabs, USA). Later, the digested insert and vector were ligated using T4 DNA ligase (M0202S, New England Biolabs, USA) followed by transformation of the ligated product into *E. coli* DH5α competent cells. The transformed colonies were screened by isolating their plasmid, subsequently double digested using *SmaI* and *XhoI* restriction enzymes. These double digested products were verified on 1% agarose gel to check for the release of insert band and the positive clones were confirmed by unidirectional Sanger sequencing.

For the generation of the V1B-gene silencing trophozoites, first stretch of 1-222 base pairs of V1B (EHI_189850) was amplified using forward (5’- gcacccgggATGGAAGCTTTCAAAATTACGCTGCTGC-3’), reverse primers (5’- gacctcgagCACTTGTACAACAGCTTTATTCCC-3’), and then cloned in pTrigger [contains 132 base pairs of the trigger gene (EHI_048600) in the pKT3M plasmid] [81] using *SmaI* and *XhoI* restriction enzymes.

### Transfection of V-ATPase subunits and generation of stable transfectants

Wild-type *E. histolytica* trophozoites were harvested in 1X phosphate buffer saline (PBS) at mid logarithmic growth phase, resuspended in incomplete cytomix buffer [10 mM K2HPO4/KH2PO4 (pH 7.6), 120 mM KCl, 0.15 mM CaCl2, 25 mM HEPES (pH 7.4), 2 mM EGTA and 5 mM MgCl2] and centrifuged at 900 x *g* for 5 min. The resultant pellets were resuspended in 0.4 mL complete cytomix buffer containing 4 mM adenosine triphosphate and 10 mM reduced glutathione, along with 80 µg of recombinant plasmid and the suspension was transferred to 4 mm electroporation cuvette (1652088 BioRad, USA). The trophozoites were then electroporated at 500 V, 500 µF, and time constant of infinity (∞) using Gene Pulser Xcell Eukaryotic System (BioRad, USA). Immediately, the contents in the electroporation cuvette were transferred into warm complete culture media (BIS-33 supplemented with 15% adult bovine serum and 2% Diamonds vitamins) without antibiotics. After 48 hours, the media was replaced with complete media and the stable transfectants were selected by gradually increasing the concentration of G418 (A1720, Sigma-Aldrich, USA) starting with 2 µg/ml to 6 µg/ml. Stable trophozoites were cultured in media containing 20 µg/ml of G418 and for inducible constructs, 30 µg/ml of tetracycline was also incorporated 48 hours before experiments.

### Western blotting

Trophozoites stably expressing V-ATPase subunits were validated for their expression levels by routine western blotting technique. Briefly, cells were harvested and lysed on ice for 30 minutes using the lysis buffer [50 mM Tris HCl (pH 7.5), 150 mM NaCl, 1 mM DTT, 1 mM PMSF, 1% Triton X-100, 10 µM E-64 (E3132, Sigma-Aldrich, USA) and protease inhibitor cocktail (PIC) (04693116001, Sigma-Aldrich, USA)]. The lysates were clarified by centrifugation at 10000 rpm for 10 minutes and protein concentration was measured using Bradford Plus protein assay (23236, Thermo Scientific, USA). Equal amounts of protein were resolved on 12% polyacrylamide gels under reducing conditions and electrophoretically transferred onto PVDF membrane (10600023, Cytiva, USA) at 90 V for 90 minutes at 4°C. Membranes were blocked in 5% bovine serum albumin (BSAV-RO, Sigma-Aldrich, USA) for 1 hour at room temperature, then probed with primary antibodies; anti-HA monoclonal (1:1000) (3724, Cell Signalling Technology, USA), anti-CS1 polyclonal (1:1000), anti-GFP (1:1000) (sc-9996, Santa Cruz Biotechnology, USA), anti-V1B (1:250) (PA5-35052, Thermo Fisher Scientific, USA), anti-actin (1:250) (MAB1501, Merck, USA) and anti-Hgl (1:100, 3F4 and 7F4). Later, the membrane was washed 3 times with 1X PBS containing 0.1% Tween 20 (PBST) for 10 minutes each, probed with HRP-conjugated anti-rabbit (1:10000) (111-035- 144, Jackson Immuno Research Laboratories, USA) or HRP-conjugated anti-mouse (1:10000) (315–035-048, Jackson Immuno Research Laboratories, USA) secondary antibody for 1 hour at room temperature. Additional 1X PBST washes were given before detecting the immunoreactive protein bands using chemiluminescence (ImageQuant™ 500, USA).

### Immunofluorescence assay

The subcellular localization of V-ATPase subunits was revealed by indirect immunofluorescence assay. Briefly, trophozoites were harvested at mid-logarithmic phase by centrifugation at 1000 x *g*, resuspended in BIS-33 and incubated on a 8-well glass slide (6040805, MP Biomedical, USA) for adherence at 37°C for 15 minutes. Cells were then fixed with 4% (w/v) paraformaldehyde (PFA) (158127, Sigma-Aldrich, USA) for 15 minutes, permeabilized with 0.1% (v/v) Triton X100 (T8787, Sigma-Aldrich, USA) for 8 minutes and blocked with 5% (v/v) FBS (26140079, Thermo Fisher Scientific, USA) for 1 hour at room temperature. Later, trophozoites were probed with primary antibodies; anti-HA monoclonal antibody (1:150) (sc-7392, Santa Cruz Biotechnology, USA), *Entamoeba* anti-V0a (1:50), or anti-PI(4,5)P2 IgM (1:50) (Z-P045, Echelon Biosciences) for 90 minutes followed by three subsequent washes with blocking solution. Secondary antibodies conjugated with Alexa Fluor 488 anti-mouse (1:500) (A11029, Invitrogen, USA), Alexa Fluor 488 anti-rabbit (1:500) (A11034, Invitrogen, USA), Alexa Fluro 568 anti-mouse (1:500) (A11031, Invitrogen, USA) or Alexa Fluor 568 Phalloidin (1:50) (A12380, Invitrogen, USA) were incubated for 1 hour at room temperature. Finally, after three additional washes with blocking solution, coverslips of thickness 1.5, 12 mm Dia (72290-04, Electron Microscopy Sciences, USA) were mounted using ProLong Diamond Antifade (P36970, Invitrogen, USA) and the slide was allowed to dry overnight at room temperature. Images were acquired using confocal microscopes.

### Colocalization quantification

Pearson’s correlation coefficient (*r*) colocalization for V-ATPase subunits and F-actin was analyzed using AIVIA 15.0.0 (aivia-software.com). Briefly, images acquired in confocal microscope were imported in AIVIA software via visualisation tool and were modified from 3D to 2D image by SwapZT tool. Trophozoites were segmented by using cellpose enhancement tool with the appropriate inputs which generates a confidence map. Further, cell tracking recipe was applied using confidence map as an input channel. Parameters (such as contrast threshold, object size, etc.) were adjusted according to the respective images. Results obtained in the spreadsheet menu were used to analyze Pearson’s correlation coefficient via advanced measurement tool.

### Cell surface biotinylation and purification

The surface associated proteins from trophozoites expressing HA-V1A were biotinylated and purified using the EZ-Link Sulfo-NHS-SS-Biotin kit (A44390, Thermo Fisher Scientific, USA) following the manufacturer’s protocol with minor modifications. Trophozoites in the mid-logarithmic growth phase (∼2 x 10^7^ trophozoites) were harvested, washed with ice-cold 1X PBS and pelleted by centrifuging at 1200 x *g* for 6 minutes. For biotin labelling, 12 mg of EZ-link Sulfo-NHS-SS-Biotin was dissolved in 10 ml ice-cold 1X PBS followed by incubation for 30 minutes at 4°C with gentle agitation on an orbital shaker to prevent endocytosis. After labelling, the cells were centrifuged and the pellet was resuspended in 10 ml 1X Tris Buffer Saline (TBS) to quench excess biotin and then pelleted again. The cell pellet was lysed in 500 µl lysis buffer (provided in the kit) supplemented with 10 µM E-64 (E3132, Sigma-Aldrich, USA) and protease inhibitor cocktail (04693116001, Sigma-Aldrich, USA) and incubated on ice for 30 minutes with sonication after every 10 minutes. The lysates were clarified by centrifuging at 13000 rpm for 10 minutes and the supernatant was incubated with NeutrAvidin Agarose beads for 1 hour at room temperature with end-over-end rotation. Prior to elution, the beads were washed 4 times with wash buffer (provided in the kit) and the biotinylated proteins were eluted in 150 µl SDS sample loading buffer containing 50 mM DTT for 1 hour at room temperature. For negative control, trophozoites were processed in the same manner as described above omitting the biotin labelling step. The elutes were subsequently resolved through SDS-PAGE and analyzed by immunoblotting.

### Treatment with various V-ATPase inhibitors

*E. histolytica* wild-type trophozoites were incubated with 0.1% DMSO (v/v) (vehicle control) (D8418, Sigma-Aldrich, USA), 50 nM concanamycin A (C9705, Sigma-Aldrich, USA) or 50 nM bafilomycin A1 (sc-201550, Santa Cruz Biotechnology, USA) in BIS-33 media for 1 hour at 35.5°C. Post treatment trophozoites were harvested by centrifugation at 1000 x *g*, resuspended in serum free media and allowed to adhere on a 8-well glass slide at 37°C for 15 minutes. The adhered cells were sequentially fixed, permeabilized, blocked and processed for indirect immunofluorescence assay as described in the “Immunofluorescence assay” methodology section, with F-actin labelling using Alexa Fluor 568 Phalloidin (A12380, Invitrogen, USA).

### Scanning Electron Microscopy

To visualize the surface topology of *E. histolytica* macropinocytic cups in the presence of V-ATPase inhibitors, scanning electron microscopy was performed. A clean 12 mm glass coverslip was placed in a 6 well culture plate. Wild-type trophozoites were treated with 0.1% DMSO, 50 nM concanamycin A and 50 nM bafilomycin A1 for 1 hour, harvested, resuspended in serum free media and incubated on a coverslip for 15 minutes at 37°C. Post incubation, media was removed, cells were rinsed with 0.1M sodium phosphate buffer (pH 7.4) for 5 minutes, fixed with 2.5% EM grade glutaraldehyde (G5882, Sigma-Aldrich) prepared in 0.1M sodium phosphate buffer (pH 7.4) and incubated overnight at 4°C. The next day, trophozoites were rinsed with 0.1M sodium phosphate buffer (pH 7.4) for 5 minutes and were dehydrated three times in ascending series of ethanol (25%, 50%, 75%, 95% and 100%) at room temperature for 5 minutes each. Coverslips were then transferred to a petri dish (lined with Kimwipes) and dried at room temperature for 72 hours. The dried samples were coated with gold using Quorum Q150T ES plus and imaged using Hitachi S-3400N scanning electron microscope (Hitachi High-Tech Corporation, Japan) at Centre for Cellular and Molecular Biology (CCMB), Hyderabad, India.

### Live cell imaging

The logarithmically grown GFP-V0d overexpressing trophozoites were harvested and resuspended in pre-warmed complete BIS-33 media, seeded on a glass-bottom 35 mm confocal dish (101350, SPL Life Sciences, Korea) and allowed to adhere for 5 min. The media was replaced with 2 mg/ml tetramethylrhodamine isothiocyanate (TRITC)-dextran (T1162, Sigma Aldrich, USA) and time-lapse images were acquired 0.863 second per frame (512 x 512) at 37 °C using Leica SP-8 confocal microscope (Leica Microsystem, Germany), equipped with HC PL APO CS2 63X oil immersion objective (NA 1.4). The 488 nm and 568 nm lasers were used for excitation of GFP and TRITC, respectively. More than five examples were collected per experiment, and one representative time lapse is provided for illustration after processed for noise reduction using LAS X software.

### Dextran uptake assay

*E. histolytica* wild-type trophozoites were serum-starved in BIS-33 media containing 0.1% DMSO (v/v) (control), 50 nM concanamycin A or 50 nM bafilomycin A1 for 1 hour at 35.5°C. Post treatment, the trophozoites were harvested, resuspended in serum-free media containing 2 mg/ml TRITC-dextran and were loaded onto a 8-well glass slide. Trophozoites were then allowed to uptake dextran for 5, 10 and 15 minutes at 37°C. Cells were fixed with 4% (w/v) PFA for 15 minutes, washed 3 times with 1X PBS and mounted with ProLong Diamond Antifade. The slides were allowed to dry overnight at room temperature prior to confocal imaging.

### Confocal microscopy

Confocal imaging was performed using LSM 900 (Carl Zeiss, Germany) or Leica SP8 confocal microscope (Leica Microsystem, Germany) equipped with 405, 488, 561, and 640 nm excitation lasers. The Plan-Apochromat 63X and HC PL APO CS2 63X oil-immersion objectives (NA 1.4) were used and all the images were acquired at 8-16-bit resolution in 512 x 512 or 1024 x 1024-pixel frame format, and scanned sequentially using standard scan speed in single en face (*xy* axes) with sections throughout the cellular *z* axis at 0.25 - 0.5 μm intervals.

### Fluorescence quantification of dextran uptake

To analyze uptake of TRITC-dextran in trophozoites, maximum intensity projection (MIP) was generated by combining all z-stacks in LAS X office and was exported in TIFF format. The MIP TIFF image was imported in Fiji ImageJ for analysis. The image type was converted from 8 bit to 16 bit. Later, threshold was adjusted and set to select only fluorescent puncta’s. Threshold was kept same for a set of images (control, bafilomycin or concanamycin treated trophozoites) throughout the analysis. Accordingly, fluorescence integrated intensities were measured using analyze tool [82]. Total integrated intensities were calculated and plotted with respect to time.

### Quantification of macropinocytic cups

The macropinocytic cups were quantified for wild type (control, bafilomycin or concanamycin treated), pEhTex-HA, HA-V1A, HA-V1B, HA-V1C and HA-V0d trophozoites irrespective of expression levels. Macropinocytic cups were manually counted for every z-stack of all the acquired confocal images avoiding the repetition of same cup in proceeding stacks. Total number of macropinocytic cups was calculated and plotted.

### Purification of recombinant GST-V1B protein

The expression and purification of GST-tagged V1B (GST-V1B) was performed by transforming the recombinant plasmid (pGEX-6P-1 bacterial expression vector harboring *Entamoeba* V1B) into chemically competent *E. coli* BL21 (DE3). Following colony screening and optimization of protein expression with various IPTG concentrations and induction time, the bacterial cultures were grown to an OD_600_ = ∼0.4 and induced with 0.5 mM IPTG (I6758, Sigma, USA) for 2 hours at 37°C. The induced cells were pelleted by centrifugation (8000 rpm for 20 minutes) and resuspended in lysis buffer (20 mM HEPES pH 6.8, 100 mM NaCl, 2 mM EDTA, 5 mM DTT and protease inhibitors) and subsequently sonicated using a probe sonicator at 35% amplitude (30 seconds ON and 30 seconds OFF), for 1 hour at 4°C. The lysates were centrifuged at 12000 rpm for 30 minutes to remove debris and the supernatant was applied to glutathione Sepharose 4B resin (17075601, Cytiva, USA) at 4°C for 16 hours to allow binding. After binding, the resin was washed 5 times with 10 bed volumes of 1X PBS after which the bound protein was eluted in glutathione elution buffer [50 mM Tris-HCl, 150 mM NaCl and 10 mM glutathione (pH 8) (Y0000517, Sigma, USA)]. The purified GST-V1B was densitometrically quantified by SDS-PAGE using BSA standards as reference.

### PA pull down assay for identification of PA-associated proteins

PA-binding proteins were captured using lipid-coated beads (P-BOPA, Echelon Biosciences, USA) in accordance with manufacturer’s guidelines. Briefly, mid-logarithmic phase wild-type trophozoites were subjected to lysis in lysis buffer (15 mM HEPES pH 7.4, 150 mM NaCl, 5 mM EDTA, 0.5% NP-40 and PIC), and clarified by centrifuging the lysate at 11000 rpm for 10 minutes at 4°C. The resulting supernatant was diluted as 1:1 with binding buffer (10 mM HEPES pH 7.4, 150 mM NaCl and 0.05% NP 40) followed by incubating with 25 µl PA-coated beads and control beads for 3 hours at 4°C under gentle rotation. The beads were washed 3 times with binding buffer, the bound proteins were eluted in SDS sample buffer by heating the lipid-bead pull-down complex at 70°C for 10 minutes. The eluted proteins were analyzed by performing LC-MS/MS based mass proteomics at Vproteomics, Noida, India (https://www.vproteomics.com/) to identify PA-associated proteins. The eluted proteins were reduced with 5 mM Tris (2-carboxyethyl) phosphine, alkylated with 50 mM iodoacetamide, and digested with trypsin at a ratio of 1:50 (trypsin:lysate) for 16 hours at 37°C. The obtained peptides were desalted on a C18 silica column to remove residual salts and dried in a vacuum concentrator. The dried peptides were resuspended in buffer A (2% acetonitrile and 0.1% formic acid) and 1 µg of the peptide was loaded on an Easy-nLC-1000 system coupled with Orbitrap Exploris mass spectrometer (Thermo Fisher Scientific, USA). The peptides were separated with Buffer B (80% acetonitrile and 0.1% formic acid) at a flow rate of 300 nl/min with 0 - 40% gradient and processed for MS analysis. The MS1 spectra were acquired in the Orbitrap (maximum injection time = 60 ms, AGC target = 300%, RF Lens = 70%, R = 60 K, mass range = 375−1500, Profile data) and MS2 spectra (maximum injection time = 60 ms, R = 15 K, AGC target 100%) was collected for the top 20 peptide ions. The raw data files were analyzed using Proteome Discoverer v2.5 (Thermo Fisher Scientific, USA) and searched against the *E. histolytica* reference database AmoebaDB (https://amoebadb.org/). For Dual Sequest and MS Amanda search, the precursor and fragment mass tolerances were set at 10 ppm and 0.02 Da, respectively. Peptide-spectrum match and protein false discovery rates were set to 1% (0.01 FDR). The identified peptides from two experimental replicates were further analyzed to obtain unique PA-binding proteins by excluding peptides in control beads samples and by avoiding the peptides with unique peptide score < 2. The comparison between overlapping and unique protein sets was identified using Venny 2.1.0 (https://bioinfogp.cnb.csic.es/tools/venny/). The bubble plot for enriched biological processes (GO_BP_Direct and UP_KW_BP) were generated in SRplot (https://www.bioinformatics.com.cn/en) using enrichment score, -log_10_(*p* value) and peptide count as input parameters. The raw mass spectrometry data have been deposited in the ProteomeXchange Consortium via the PRIDE [83] partner repository under the dataset identifiers PXD075324 (PA-interactome).

### Actin co-sedimentation assay

To test whether the *E. histolytica* V1B subunit can bind to G-actin and polymerize G-actin to F-actin, the Actin Polymerization Biochem Kit (BK001, Cytoskeleton, USA) was used as per the manufacturer’s instructions. For *in vitro* binding reactions, F-actin stock was prepared by resuspending 250 µg of rabbit muscle actin in general actin buffer and incubated for 30 minutes on ice. To initiate F-actin formation, 20 µl actin polymerization buffer was added and incubated for 60 minutes at room temperature. Before setting the reaction, both bovine serum albumin (BSA) and recombinant protein (GST-V1B) were clarified using ultracentrifugation equipped with TLA-100 rotor (Optima XPN-100, Beckman Coulter, USA) at 150000 x *g* for 1 hour at 4°C to remove any aggregates. Binding reactions were prepared by incubating 10 µl of the purified GST-V1B protein (15 µM) with 40 µl F-actin stock at room temperature for 30 minutes. For *in vitro* polymerization reactions, rabbit muscle G-actin (21 µM) was resuspended in general actin buffer supplemented with 0.2 mM ATP and incubated on ice for 1 hour to prepare monomeric G-actin stock. Before setting the reaction, both BSA and the recombinant protein GST-V1B were clarified using ultracentrifugation equipped with TLA-100 rotor (Optima XPN-100, Beckman Coulter, USA) at 150000 x *g* for 1 hour at 4°C to remove any aggregates. Polymerization reactions were prepared by incubating 10µl of the purified GST-V1B protein (15 µM) with 40 µl G-actin stock at room temperature for 30 minutes. For both binding and polymerization reactions, incubation of α-actinin (positive control) and BSA (negative control) were set in parallel with the test protein. The co-sedimentation of GST-V1B with polymerized F-actin (binding and polymerization) was analyzed by resolving the isolated supernatant and pellet fractions by SDS-PAGE and stained with Coomassie Brilliant Blue G-250 (1610406, BioRad, USA). The respective band intensities of proteins in supernatant and pellet fractions were quantified using ImageJ software.

### Pyrene-labelled actin polymerization assay

For quantitative assessment of GST-V1B induced actin polymerization, the actin polymerization Biochem Kit (BK003, Cytoskeleton, USA) was used as per the manufacturer’s instructions. Briefly, pyrene labelled G-actin (0.4 mg/mL) was prepared in general actin buffer supplemented with 0.2 mM ATP and equilibrated on ice for 1 hour. The prepared G-actin was centrifuged at 14000 rpm for 30 minutes at 4°C to remove aggregates. Baseline curve was obtained by recording the fluorescence 3 - 5 µM pyrene-labelled G-actin (200 µl/condition) to confirm the equal distribution of pyrene actin in every condition. Polymerization reactions were set by adding various concentrations of GST-V1B (0.625, 1.25, 2.5 and 5 µM) and GST (negative control) in the same wells containing pyrene labelled G-actin and actin polymerization buffer in black 96-well plate. To evaluate the effect of PA on GST-V1B-mediated actin polymerization, PA (P9511, Sigma Aldrich, USA) was utilized at different concentrations (2.5, 5, 10 and 20 µM) along with GST-V1B (0.625 µM) under the same experimental setup as described above. Fluorescence readings (excitation 350 nm, emission 420 nm) were recorded kinetically at room temperature for every 1 minute over a period of 60 minutes using microplate reader (SpectraMax iD5, Molecular devices, USA). Polymerization curves were obtained by plotting raw fluorescence values.

### Lipid overlay assay

PIP Strip membranes (P23750, Thermo Fisher Scientific, USA) spotted with various phosphoinositides, were used to assess the lipid binding specificity of GST-V1B. PIP strips were equilibrated in 1X TBST (100 mM tris, 0.15 M sodium chloride and 0.5% Tween 20^®^) containing 3% (w/v) fatty acid-free bovine serum albumin (BSA) (SKU:02199899-CF, MP Biomedical, USA) for 1 hour at room temperature. The equilibrated membranes were probed with purified GST and GST-V1B (0.7 µg/ml) diluted in the lipid-binding solution (TBS-T + 3% fatty acid–free BSA) and incubated overnight at 4°C on an orbital shaker. After incubation, the membranes were washed 3 times with binding solution and probed with an anti-GST primary antibody (1:500) (B-14:sc-138, Santa Cruz, USA) followed by horseradish peroxidase-conjugated anti-rabbit secondary antibody (1:10000) (111-035-144, Jackson Immuno Research Laboratories, USA). For detection, the strips were washed three times and the bound lipid-proteins complexes were visualized using chemiluminescence (ImageQuant™ 500, USA). The individual spot intensities were quantified using ImageJ. **Statistical analysis**

Statistical significance of the graphs corresponding to macropinocytic cup number, their association with V-ATPase, dextran uptake and lipid binding assay were calculated using Student’s *t*-test in GraphPad Prism 9. The sample size and obtained *p*-values are indicated in the respective figure legends.

## Data availability

All relevant data are within the manuscript and its supplementary information files.

## Author Contributions

**Bhagyashree Chordiya** and **Navyaka Padavala**: Formal analysis; Investigation; Visualization; Methodology. **Amisha Sharma and Kadambini Pradhan**: Formal analysis; Investigation; Methodology: **Kuldeep Verma**: Conceptualization; Funding acquisition; Visualization; Project administration; Resources; Supervision; Writing-review and editing.

## Acknowledgements

We are thankful to Prof. Tomoyoshi Nozaki (Graduate School of Medicine, University of Tokyo, Japan) and Prof. William A. Petri Jr. (University of Virginia, USA), for generously providing the anti-CS1, anti-Hgl antibodies, respectively. We thank to Dr. Rupinder Kaur (BRIC-CDFD, Hyderabad, India) and Prof. Sunando Datta (IISER, Bhopal, India), for helpful discussion and feedback on the manuscript. We thank Sanjana Parida (Dissertation trainee) for excellent technical help in protein expression optimization. We also thank Sandeep Srivastava (Electron Microscopy Facility, CSIR-CCMB, Hyderabad, India) for helping in SEM imaging. We thank all the members of the KV laboratory for helpful discussions.

## References

1. Tu, H., Wang, H., and Cai, H. (2025). Macropinocytosis: Molecular mechanisms and regulation. Curr Opin Cell Biol 95, 102563.

2. Bohdanowicz, M., Schlam, D., Hermansson, M., Rizzuti, D., Fairn, G.D., Ueyama, T., Somerharju, P., Du, G., and Grinstein, S. (2013). Phosphatidic acid is required for the constitutive ruffling and macropinocytosis of phagocytes. Mol Biol Cell 24, 1700–1712, S1701-1707.

3. Bohdanowicz, M., Cosio, G., Backer, J.M., and Grinstein, S. (2010). Class I and class III phosphoinositide 3-kinases are required for actin polymerization that propels phagosomes. J Cell Biol 191, 999–1012.

4. Swanson, J.A. (2008). Shaping cups into phagosomes and macropinosomes. Nat Rev Mol Cell Biol 9, 639–649.

5. Buckley, C.M., and King, J.S. (2017). Drinking problems: mechanisms of macropinosome formation and maturation. FEBS J 284, 3778–3790.

6. King, J.S., and Kay, R.R. (2019). The origins and evolution of macropinocytosis. Philos Trans R Soc Lond B Biol Sci 374, 20180158.

7. Hacker, U., Albrecht, R., and Maniak, M. (1997). Fluid-phase uptake by macropinocytosis in Dictyostelium. J Cell Sci 110 (Pt 2), 105–112.

8. Khan, N.A. (2006). Acanthamoeba: biology and increasing importance in human health. FEMS Microbiol Rev 30, 564–595.

9. Kawashima, A., Yanagawa, Y., Shimogawara, R., Yagita, K., Gatanaga, H., and Watanabe, K. (2023). Amebiasis as a sexually transmitted infection: A re-emerging health problem in developed countries. Glob Health Med 5, 319–327.

10. Shirley, D.T., Farr, L., Watanabe, K., and Moonah, S. (2018). A Review of the Global Burden, New Diagnostics, and Current Therapeutics for Amebiasis. Open Forum Infect Dis 5, ofy161.

11. Lozano, R., Naghavi, M., Foreman, K., Lim, S., Shibuya, K., Aboyans, V., Abraham, J., Adair, T., Aggarwal, R., Ahn, S.Y., et al. (2012). Global and regional mortality from 235 causes of death for 20 age groups in 1990 and 2010: a systematic analysis for the Global Burden of Disease Study 2010. Lancet 380, 2095–2128.

12. Group, I.P.C. (2024). Global burden associated with 85 pathogens in 2019: a systematic analysis for the Global Burden of Disease Study 2019. Lancet Infect Dis 24, 868–895.

13. Miller, H.W., Suleiman, R.L., and Ralston, K.S. (2019). Trogocytosis by Entamoeba histolytica Mediates Acquisition and Display of Human Cell Membrane Proteins and Evasion of Lysis by Human Serum. mBio 10.

14. Guillen, N. (2021). Signals and signal transduction pathways in Entamoeba histolytica during the life cycle and when interacting with bacteria or human cells. Mol Microbiol 115, 901–915.

15. Uddin, M.J., Leslie, J.L., and Petri, W.A., Jr. (2021). Host Protective Mechanisms to Intestinal Amebiasis. Trends Parasitol 37, 165–175.

16. Meza, I., and Clarke, M. (2004). Dynamics of endocytic traffic of Entamoeba histolytica revealed by confocal microscopy and flow cytometry. Cell Motil Cytoskeleton 59, 215–226.

17. Quinn, S.E., Huang, L., Kerkvliet, J.G., Swanson, J.A., Smith, S., Hoppe, A.D., Anderson, R.B., Thiex, N.W., and Scott, B.L. (2021). The structural dynamics of macropinosome formation and PI3-kinase- mediated sealing revealed by lattice light sheet microscopy. Nat Commun 12, 4838.

18. Apte, A., Manich, M., Labruyere, E., and Datta, S. (2022). PI Kinase-EhGEF2-EhRho5 axis contributes to LPA stimulated macropinocytosis in Entamoeba histolytica. PLoS Pathog 18, e1010550.

19. Shimoyama, M., Nakada-Tsukui, K., and Nozaki, T. (2024). EhRacM differentially regulates macropinocytosis and motility in the enteric protozoan parasite Entamoeba histolytica. PLoS Pathog 20, e1012364.

20. Kadri, S., Nakada-Tsukui, K., Watanabe, N., Jeelani, G., and Nozaki, T. (2022). PTEN differentially regulates endocytosis, migration, and proliferation in the enteric protozoan parasite Entamoeba histolytica. PLoS Pathog 18, e1010147.

21. Gilmartin, A.A., Ralston, K.S., and Petri, W.A., Jr. (2017). Inhibition of Amebic Lysosomal Acidification Blocks Amebic Trogocytosis and Cell Killing. mBio 8.

22. Mitra, B.N., Yasuda, T., Kobayashi, S., Saito-Nakano, Y., and Nozaki, T. (2005). Differences in morphology of phagosomes and kinetics of acidification and degradation in phagosomes between the pathogenic Entamoeba histolytica and the non-pathogenic Entamoeba dispar. Cell Motil Cytoskeleton 62, 84–99.

23. Maxson, M.E., and Grinstein, S. (2014). The vacuolar-type H(+)-ATPase at a glance - more than a proton pump. J Cell Sci 127, 4987–4993.

24. Breton, S., and Brown, D. (2013). Regulation of luminal acidification by the V-ATPase. Physiology (Bethesda) 28, 318–329.

25. Wang, R., Long, T., Hassan, A., Wang, J., Sun, Y., Xie, X.S., and Li, X. (2020). Cryo-EM structures of intact V-ATPase from bovine brain. Nat Commun 11, 3921.

26. Zhao, J., Benlekbir, S., and Rubinstein, J.L. (2015). Electron cryomicroscopy observation of rotational states in a eukaryotic V-ATPase. Nature 521, 241–245.

27. Roh, S.H., Stam, N.J., Hryc, C.F., Couoh-Cardel, S., Pintilie, G., Chiu, W., and Wilkens, S. (2018). The 3.5- A CryoEM Structure of Nanodisc-Reconstituted Yeast Vacuolar ATPase V(o) Proton Channel. Mol Cell 69, 993–1004 e1003.

28. Mazhab-Jafari, M.T., Rohou, A., Schmidt, C., Bueler, S.A., Benlekbir, S., Robinson, C.V., and Rubinstein, J.L. (2016). Atomic model for the membrane-embedded V(O) motor of a eukaryotic V-ATPase. Nature 539, 118–122.

29. Jaskolka, M.C., Tarsio, M., Smardon, A.M., Khan, M.M., and Kane, P.M. (2021). Defining steps in RAVE- catalyzed V-ATPase assembly using purified RAVE and V-ATPase subcomplexes. J Biol Chem 296, 100703.

30. Holliday, L.S., Lu, M., Lee, B.S., Nelson, R.D., Solivan, S., Zhang, L., and Gluck, S.L. (2000). The amino- terminal domain of the B subunit of vacuolar H+-ATPase contains a filamentous actin binding site. J Biol Chem 275, 32331–32337.

31. Vitavska, O., Wieczorek, H., and Merzendorfer, H. (2003). A novel role for subunit C in mediating binding of the H+-V-ATPase to the actin cytoskeleton. J Biol Chem 278, 18499–18505.

32. Ma, B., Qian, D., Nan, Q., Tan, C., An, L., and Xiang, Y. (2012). Arabidopsis vacuolar H+-ATPase (V- ATPase) B subunits are involved in actin cytoskeleton remodeling via binding to, bundling, and stabilizing F-actin. J Biol Chem 287, 19008–19017.

33. Vitavska, O., Merzendorfer, H., and Wieczorek, H. (2005). The V-ATPase subunit C binds to polymeric F-actin as well as to monomeric G-actin and induces cross-linking of actin filaments. J Biol Chem 280, 1070–1076.

34. Petzoldt, A.G., Gleixner, E.M., Fumagalli, A., Vaccari, T., and Simons, M. (2013). Elevated expression of the V-ATPase C subunit triggers JNK-dependent cell invasion and overgrowth in a Drosophila epithelium. Dis Model Mech 6, 689–700.

35. Carnell, M., Zech, T., Calaminus, S.D., Ura, S., Hagedorn, M., Johnston, S.A., May, R.C., Soldati, T., Machesky, L.M., and Insall, R.H. (2011). Actin polymerization driven by WASH causes V-ATPase retrieval and vesicle neutralization before exocytosis. J Cell Biol 193, 831–839.

36. Serra-Peinado, C., Sicart, A., Llopis, J., and Egea, G. (2016). Actin Filaments Are Involved in the Coupling of V0-V1 Domains of Vacuolar H+-ATPase at the Golgi Complex. J Biol Chem 291, 7286–7299.

37. Ramirez, C., Hauser, A.D., Vucic, E.A., and Bar-Sagi, D. (2019). Plasma membrane V-ATPase controls oncogenic RAS-induced macropinocytosis. Nature 576, 477–481.

38. Melendez-Hernandez, M.G., Barrios, M.L., Orozco, E., and Luna-Arias, J.P. (2008). The vacuolar ATPase from Entamoeba histolytica: molecular cloning of the gene encoding for the B subunit and subcellular localization of the protein. BMC Microbiol 8, 235.

39. Yi, Y., and Samuelson, J. (1994). Primary structure of the Entamoeba histolytica gene (Ehvma1) encoding the catalytic peptide of a putative vacuolar membrane proton-transporting ATPase (V-ATPase). Mol Biochem Parasitol 66, 165–169.

40. Cotter, K., Capecci, J., Sennoune, S., Huss, M., Maier, M., Martinez-Zaguilan, R., and Forgac, M. (2015). Activity of plasma membrane V-ATPases is critical for the invasion of MDA-MB231 breast cancer cells. J Biol Chem 290, 3680–3692.

41. Sumner, J.P., Dow, J.A., Earley, F.G., Klein, U., Jager, D., and Wieczorek, H. (1995). Regulation of plasma membrane V-ATPase activity by dissociation of peripheral subunits. J Biol Chem 270, 5649–5653.

42. Saw, N.M., Kang, S.Y., Parsaud, L., Han, G.A., Jiang, T., Grzegorczyk, K., Surkont, M., Sun-Wada, G.H., Wada, Y., Li, L., et al. (2011). Vacuolar H(+)-ATPase subunits Voa1 and Voa2 cooperatively regulate secretory vesicle acidification, transmitter uptake, and storage. Mol Biol Cell 22, 3394–3409.

43. Posor, Y., Jang, W., and Haucke, V. (2022). Phosphoinositides as membrane organizers. Nat Rev Mol Cell Biol 23, 797–816.

44. Padavala, N., and Verma, K. (2025). Glutamic acid-lysine (EK) rich motif of RabD2 self-associates and regulates adhesion through multivesicular bodies in Entamoeba histolytica. BMC Biol 23, 308.

45. Biller, L., Matthiesen, J., Kuhne, V., Lotter, H., Handal, G., Nozaki, T., Saito-Nakano, Y., Schumann, M., Roeder, T., Tannich, E., et al. (2014). The cell surface proteome of Entamoeba histolytica. Mol Cell Proteomics 13, 132–144.

46. Petri, W.A., Jr., Chapman, M.D., Snodgrass, T., Mann, B.J., Broman, J., and Ravdin, J.I. (1989). Subunit structure of the galactose and N-acetyl-D-galactosamine-inhibitable adherence lectin of Entamoeba histolytica. J Biol Chem 264, 3007–3012.

47. Sava, I., Davis, L.J., Gray, S.R., Bright, N.A., and Luzio, J.P. (2024). Reversible assembly and disassembly of V-ATPase during the lysosome regeneration cycle. Mol Biol Cell 35, ar63.

48. Nakada-Tsukui, K., Saito-Nakano, Y., Ali, V., and Nozaki, T. (2005). A retromerlike complex is a novel Rab7 effector that is involved in the transport of the virulence factor cysteine protease in the enteric protozoan parasite Entamoeba histolytica. Mol Biol Cell 16, 5294–5303.

49. Verma, K., and Datta, S. (2017). The Monomeric GTPase Rab35 Regulates Phagocytic Cup Formation and Phagosomal Maturation in Entamoeba histolytica. J Biol Chem 292, 4960–4975.

50. Manich, M., Hernandez-Cuevas, N., Ospina-Villa, J.D., Syan, S., Marchat, L.A., Olivo-Marin, J.C., and Guillen, N. (2018). Morphodynamics of the Actin-Rich Cytoskeleton in Entamoeba histolytica. Front Cell Infect Microbiol 8, 179.

51. Huss, M., Ingenhorst, G., Konig, S., Gassel, M., Drose, S., Zeeck, A., Altendorf, K., and Wieczorek, H. (2002). Concanamycin A, the specific inhibitor of V-ATPases, binds to the V(o) subunit c. J Biol Chem 277, 40544–40548.

52. Wang, R., Wang, J., Hassan, A., Lee, C.H., Xie, X.S., and Li, X. (2021). Molecular basis of V-ATPase inhibition by bafilomycin A1. Nat Commun 12, 1782.

53. Mercanti, V., Charette, S.J., Bennett, N., Ryckewaert, J.J., Letourneur, F., and Cosson, P. (2006). Selective membrane exclusion in phagocytic and macropinocytic cups. J Cell Sci 119, 4079–4087.

54. Buckley, C.M., Pots, H., Gueho, A., Vines, J.H., Munn, C.J., Phillips, B.A., Gilsbach, B., Traynor, D., Nikolaev, A., Soldati, T., et al. (2020). Coordinated Ras and Rac Activity Shapes Macropinocytic Cups and Enables Phagocytosis of Geometrically Diverse Bacteria. Curr Biol 30, 2912–2926 e2915.

55. Young, B.P., Shin, J.J., Orij, R., Chao, J.T., Li, S.C., Guan, X.L., Khong, A., Jan, E., Wenk, M.R., Prinz, W.A., et al. (2010). Phosphatidic acid is a pH biosensor that links membrane biogenesis to metabolism. Science 329, 1085–1088.

56. Ha, K.S., and Exton, J.H. (1993). Activation of actin polymerization by phosphatidic acid derived from phosphatidylcholine in IIC9 fibroblasts. J Cell Biol 123, 1789–1796.

57. Zhou, H., Huo, Y., Yang, N., and Wei, T. (2024). Phosphatidic acid: from biophysical properties to diverse functions. FEBS J 291, 1870–1885.

58. Fang, Y., Vilella-Bach, M., Bachmann, R., Flanigan, A., and Chen, J. (2001). Phosphatidic acid-mediated mitogenic activation of mTOR signaling. Science 294, 1942–1945.

59. Kooijman, E.E., Chupin, V., de Kruijff, B., and Burger, K.N. (2003). Modulation of membrane curvature by phosphatidic acid and lysophosphatidic acid. Traffic 4, 162–174.

60. Huang, S., Gao, L., Blanchoin, L., and Staiger, C.J. (2006). Heterodimeric capping protein from Arabidopsis is regulated by phosphatidic acid. Mol Biol Cell 17, 1946–1958.

61. Lohden-Bendinger, U., and Bakker-Grunwald, T. (1990). Evidence for a vacuolar-type proton ATPase in Entamoeba histolytica. Z Naturforsch C J Biosci 45, 229–232.

62. Gluck, S. (1992). V-ATPases of the plasma membrane. J Exp Biol 172, 29–37.

63. Rojas, J.D., Sennoune, S.R., Maiti, D., Bakunts, K., Reuveni, M., Sanka, S.C., Martinez, G.M., Seftor, E.A., Meininger, C.J., Wu, G., et al. (2006). Vacuolar-type H+-ATPases at the plasma membrane regulate pH and cell migration in microvascular endothelial cells. Am J Physiol Heart Circ Physiol 291, H1147–1157.

64. Proctor, E.M., and Gregory, M.A. (1972). The observation of a surface active lysosome in the trophozoites of Entamoeba histolytica from the human colon. Ann Trop Med Parasitol 66, 339–342.

65. Eaton, R.D., Meerovitch, E., and Costerton, J.W. (1969). A surface-active lysosome in Entamoeba histolytica. Trans R Soc Trop Med Hyg 63, 678–680.

66. Toyomura, T., Murata, Y., Yamamoto, A., Oka, T., Sun-Wada, G.H., Wada, Y., and Futai, M. (2003). From lysosomes to the plasma membrane: localization of vacuolar-type H+ -ATPase with the a3 isoform during osteoclast differentiation. J Biol Chem 278, 22023–22030.

67. Pastor-Soler, N.M., Hallows, K.R., Smolak, C., Gong, F., Brown, D., and Breton, S. (2008). Alkaline pH- and cAMP-induced V-ATPase membrane accumulation is mediated by protein kinase A in epididymal clear cells. Am J Physiol Cell Physiol 294, C488–494.

68. Cotter, K., Liberman, R., Sun-Wada, G., Wada, Y., Sgroi, D., Naber, S., Brown, D., Breton, S., and Forgac, M. (2016). The a3 isoform of subunit a of the vacuolar ATPase localizes to the plasma membrane of invasive breast tumor cells and is overexpressed in human breast cancer. Oncotarget 7, 46142–46157.

69. Sautin, Y.Y., Lu, M., Gaugler, A., Zhang, L., and Gluck, S.L. (2005). Phosphatidylinositol 3-kinase- mediated effects of glucose on vacuolar H+-ATPase assembly, translocation, and acidification of intracellular compartments in renal epithelial cells. Mol Cell Biol 25, 575–589.

70. McGuire, C.M., and Forgac, M. (2018). Glucose starvation increases V-ATPase assembly and activity in mammalian cells through AMP kinase and phosphatidylinositide 3-kinase/Akt signaling. J Biol Chem 293, 9113–9123.

71. Koushik, A.B., Welter, B.H., Rock, M.L., and Temesvari, L.A. (2014). A genomewide overexpression screen identifies genes involved in the phosphatidylinositol 3-kinase pathway in the human protozoan parasite Entamoeba histolytica. Eukaryot Cell 13, 401–411.

72. Lee, Y.A., Kim, K.A., Min, A., and Shin, M.H. (2014). Amoebic PI3K and PKC is required for Jurkat T cell death induced by Entamoeba histolytica. Korean J Parasitol 52, 355–365.

73. Byekova, Y.A., Powell, R.R., Welter, B.H., and Temesvari, L.A. (2010). Localization of phosphatidylinositol (3,4,5)-trisphosphate to phagosomes in entamoeba histolytica achieved using glutathione S-transferase- and green fluorescent protein-tagged lipid biosensors. Infect Immun 78, 125–137.

74. Bowman, E.J., Graham, L.A., Stevens, T.H., and Bowman, B.J. (2004). The bafilomycin/concanamycin binding site in subunit c of the V-ATPases from Neurospora crassa and Saccharomyces cerevisiae. J Biol Chem 279, 33131–33138.

75. Roach, A.N., Wang, Z., Wu, P., Zhang, F., Chan, R.B., Yonekubo, Y., Di Paolo, G., Gorfe, A.A., and Du, G. (2012). Phosphatidic acid regulation of PIPKI is critical for actin cytoskeletal reorganization. J Lipid Res 53, 2598–2609.

76. Behnen, M., Murk, K., Kursula, P., Cappallo-Obermann, H., Rothkegel, M., Kierszenbaum, A.L., and Kirchhoff, C. (2009). Testis-expressed profilins 3 and 4 show distinct functional characteristics and localize in the acroplaxome-manchette complex in spermatids. BMC Cell Biol 10, 34.

77. Antonescu, C.N., Danuser, G., and Schmid, S.L. (2010). Phosphatidic acid plays a regulatory role in clathrin-mediated endocytosis. Mol Biol Cell 21, 2944–2952.

78. Sherr, G.L., LaMassa, N., Li, E., Phillips, G., and Shen, C.H. (2017). Pah1p negatively regulates the expression of V-ATPase genes as well as vacuolar acidification. Biochem Biophys Res Commun 491, 693–700.

79. Wang, Q., Wolf, A., Ozkan, S., Richert, L., Mely, Y., Chasserot-Golaz, S., Ory, S., Gasman, S., and Vitale, N. (2023). V-ATPase modulates exocytosis in neuroendocrine cells through the activation of the ARNO-Arf6-PLD pathway and the synthesis of phosphatidic acid. Front Mol Biosci 10, 1163545.

80. Mitra, C., Winkley, S., and Kane, P.M. (2023). Human V-ATPase a-subunit isoforms bind specifically to distinct phosphoinositide phospholipids. J Biol Chem 299, 105473.

81. Morf, L., Pearson, R.J., Wang, A.S., and Singh, U. (2013). Robust gene silencing mediated by antisense small RNAs in the pathogenic protist Entamoeba histolytica. Nucleic Acids Res 41, 9424–9437.

82. Schindelin, J., Arganda-Carreras, I., Frise, E., Kaynig, V., Longair, M., Pietzsch, T., Preibisch, S., Rueden, C., Saalfeld, S., Schmid, B., et al. (2012). Fiji: an open-source platform for biological-image analysis. Nat Methods 9, 676–682.

83. Perez-Riverol, Y., Bandla, C., Kundu, D.J., Kamatchinathan, S., Bai, J., Hewapathirana, S., John, N.S., Prakash, A., Walzer, M., Wang, S., et al. (2025). The PRIDE database at 20 years: 2025 update. Nucleic Acids Res 53, D543–D553.

