## Supplementary material for "Evolutionary divergence V-ATPase function in macropinocytic cup remodeling": Supplementary Information.docx

**Supplementary Methods**

**Evolutionary conservation of V-ATPase subunits**

To study the evolutionary conservation of *E. histolytica* V-ATPase subunits across diverse eukaryotes, *in silico* analysis was performed. The protein sequences of *E. histolytica* V-ATPase subunits were retrieved from AmoebaDB (amoebadb.org) and orthologous sequences of *Homo sapiens, Arabidopsis thaliana, Saccharomyces cerevisiae, Plasmodium falciparum, Dictyostelium discoideum, Drosophila melanogaster, Caenorhabditis elegans, Xenopus laevis* were retrieved from UniProtKB (uniprot.org). Pairwise Blastp analysis was performed for each V-ATPase subunit using *E. histolytica* sequences as query against the listed orthologs to yield the percent identity (Table S1).

***Entamoeba* V0a subunit antibody generation and validation**

To assess the localization of V0a subunit and to monitor the recruitment of V-ATPase pump, a synthetic peptide antigen was designed against the N-terminal region (74-102 amino acids) of *E. histolytica* V0a subunit (EHI_074020, predicted mass ~ 92.6 kDa). Based on the predicted antigenic determinants, two N-terminal peptide epitopes (RNGEFQSFEKESTEC and KEEERKDMNGLKFRC) were concatenated into a 30-mer single peptide, which was synthesized by GeneScript (USA). For immunization, an initial dose of 500 µg peptide resuspended in 500 µl 1X PBS, emulsified with Freund’s adjuvant was injected subcutaneously into an 8-week old New-Zealand White rabbit. Booster doses with increased amount of peptide i.e., 750 µg prepared in 500 µl 1X PBS was administered on days 50, 70, 90 and 110. Pre-immune serum was collected prior to the first injection and test bleeds were obtained after the final booster. For indirect immunofluorescence assay, dilutions along a wide range 1:25, 1:50 and 1:100 were tested with the anti-V0a antibody. The 1:50 dilution reflected specific signal with low background and therefore used in all the experiments. To further confirm the specificity of antibody, peptide competition assay was performed. The anti-V0a serum was pre-incubated with a 20-fold excess synthetic peptide (25 µg) for 90 minutes at room temperature to allow the saturation of antibody binding sites with the free peptide. This peptide-antibody mixture was used as a primary antibody in standard immunofluorescence assay. In parallel, BSA was used as a control and images were acquired using confocal microscope.

Western blotting was performed to assess the anti-V0a reactivity against the whole cell lysate of wild-type trophozoites using 1:20 and 1:50 dilutions; however, the antibody was confirmed to be unsuitable for western blotting. To further validate this antibody, membrane fraction of the trophozoites were isolated. Approximate 2x10^7^ *E. histolytica* trophozoites were harvested by centrifugation at 1200 x g for 6 minutes. The resulting cell pellet was resuspended in 3 ml of lysis buffer containing 10 mM HEPES (pH 7.5), 1.5 mM MgCl₂, 10 mM KCl, 0.5 mM DTT, 0.2% NP-40, 0.1 M PMSF, 10 µM E64, and protease inhibitor cocktail. The suspension was incubated on ice for 15 minutes and subsequently centrifuged at 3000 x g for 10 minutes. The pellet obtained representing the nuclear fraction. The supernatant was transferred to a fresh microcentrifuge tube and was subjected to ultra-centrifugation at 100,000 x g for 30 minutes at 4°C using a 90Ti rotor in 2 ml tubes. The resulting pellet, corresponding to the membrane fraction, was further resuspended in 100 µl of lysis buffer and sonicated (5 sec ON/OFF cycle for 1 min). Bradford assay was used to determine the protein concentration. Total 50 µg concentration of this membrane fraction was subjected to SDS-PAGE and probed with V0a antibody (1:50 dilution). **Live immunofluorescence assay**

To confirm whether the generated V0a antibody binds to the extracellular epitopes of V0a subunit in trophozoites, live immunofluorescence staining was performed as described with few modifications (Biller et al., 2014). Amoebic wild-type trophozoites were harvested in mid-logarithmic phase at 1200 x *g* for 5 minutes at 4°C and resuspended in 2 ml of blocking buffer (50% v/v BIS-33 + 50% v/v 1X PBS containing 20% FBS+ 20 µM E64), followed by incubation for 10 minutes at 4°C. The trophozoites were sedimented again and resuspended in 400 µl of primary antibody; anti-V0a (1:50) for 20 minutes at 4°C. Post incubation, trophozoites were washed with the blocking buffer for 3 times. The trophozoites were incubated with 400 µl of secondary antibody conjugated with Alexa fluor 488 anti-rabbit (1:500) for 20 minutes at 4°C followed by 3 washes with blocking buffer and one wash with 1X PBS. Trophozoites were then fixed with 4% PFA for 10 minutes at 4°C. The trophozoites were then loaded on confocal dish with glass bottom and images were acquired using confocal microscope.

**LysoTracker red staining**

The HA-V1A and HA-V0d expressing trophozoites were incubated with 2 µM LysoTracker Red DND-99 (L7528, Invitrogen, USA) in BIS-33 medium for 1 hour at 35.5°C as reported previously with minor modification (Verma and Datta, 2017). Post incubation, trophozoites were harvested by centrifugation at 1000 x *g* and loaded on a 8-well glass slide for adherence for 15 minutes at 37°C. Cells were then fixed with 4% PFA for 15 minutes at 37°C, permeabilized with 0.1% (v/v) Triton X100 for 8 minutes and blocked with 5% (v/v) FBS for 1 hour at room temperature. Later, trophozoites were probed with anti-HA monoclonal antibody (1:150) for 90 minutes at room temperature, followed by three subsequent washes with blocking solution. Further, probed with secondary antibody conjugated with Alexa Fluor-488 anti-mouse (1:500) for 1 hour at room temperature. Finally, after three washes with blocking solution, coverslips were mounted using 5 µL ProLong Diamond Antifade and the slide was allowed to dry overnight at room temperature. Images were acquired using confocal microscope.

**Quantification of Lysotracker red compartments**

The acidic compartments were quantified for HA-V1A and HA-V0d expressing trophozoites. The lysotracker positive compartments were manually quantified by counting lysotracker puncta in every z-stack eliminating the repeated puncta in proceeding stacks. The acidic compartments associated with V-ATPase were also quantified simultaneously.

**Supplementary Figure legends**

**Figure S1. Cloning of V-ATPase subunits of *E. histolytica***

The putative Entamoeba V1A (EHI_043010), V1B (EHI_189850), V1C (EHI_092600) and V0d (EHI_106350) genes were amplified by PCR using gene specific forward and reverse primers from cDNA pool of *E. histolytica*.

**Figure S2. Validation of *E. histolytica* V-ATPase V0a antibody**

A. Immunofluorescence assay was performed to validate the specificity of anti-V0a antibody. *E. histolytica* wild-type trophozoites were fixed, permeabilized and probed with pre-immune sera (collected from the rabbit prior to injecting the V0a peptide) (1:50) and anti-V0a (1:50) polyclonal sera, followed by secondary antibody Alexa fluor-488. The *white* square object represents a zoomed panel. Scale bars, 10 µm

B. Wild-type *E. histolytica* trophozoites were fixed and probed with anti-V0a polyclonal sera (1:50, upper lane), followed by secondary antibody Alexa fluor-488. The confocal images show the fluorescence signals specific to cell surface. The *white* square object represents a zoomed panel. Scale bars, 10 µm

C. Live cell immunofluorescence assay for V0a. Live wild-type trophozoites were incubated with anti-V0a at 4°C, then probed with secondary antibody Alexa fluor-488. Finally, trophozoites were fixed and imaged using confocal microscopy. The *white* square object represents a zoomed panel. Scale bars, 10 µm

D. *E. histolytica* wild-type trophozoites were fixed, permeabilized and incubated with the mixture of anti-V0a (1:50) and0.5 µg/µl V0a peptide (upper panel) and anti-V0a (1:50) with 0.5 µg/µl BSA (lower panel), followed by secondary antibody Alexa fluor-488. The confocal images showed that, in the presence of the V0a peptide, the anti-V0a antibody failed to produce a fluorescence signal, whereas incubation with BSA did not affect antibody binding. This confirms that the produced antibody specifically recognizes the V0a subunit. The *white* square object represents a zoomed panel. Scale bars, 10 µm

E. The membrane fractions (50 µg) of wild-type trophozoites were resolved and was subjected to immunoblotting with *Entamoeba* V0a antibody. The *red* arrowhead indicates the V0a predicted size around 92 kDa.

**Figure S3. Association of V-ATPase subunits with LysoTracker-Red positive compartments**

HA-V1A and HA-V0d expressing trophozoites were incubated with 2 µM LysoTracker for 1 hour in BIS-33 media at 37°C. Subsequently, trophozoites were fixed and processed with anti-HA antibody. *White* arrowheads indicate HA-V1A (upper panel) and HA-V0d (lower panel) associated with LysoTracker-positive compartments. While *white* arrows indicate the V-ATPase compartment negative for LysoTracker. Scale bars, 10 µm

**Figure S4. Representative images for colocalization studies**

Amoebic trophozoites stably expressing the V-ATPase subunits (HA-V1A, HA-V1B, HA-V1C, and HA-V0d), were incubated for 15 min at 37°C under steady state condition. Post incubation, trophozoites were processed for immunofluorescence using anti-HA and phalloidin. The *Z*-stack of confocal images were acquired and colocalization between each V-ATPase subunit and F-actin was quantified using Aivia 15 software. Scale bars, 5 µm

**Figure S5. Rabbit anti-V1B antibody (raised against human V1B epitope) recognizes *E. histolytica* endogenous V1B.**

**A**. Sequence alignment of human V1B1 and *E. histolytica* V1B subunits. The alignment highlights conserved epitope residues between the two proteins. The solid black line marks the corresponding residues of the antibody epitope region. High conservation (>90% identity) supports the rationale for using the human V1B1 antibody to detect *E. histolytica* V1B.

**B, C.** Whole lysate (40 µg) of HepG2 cells and vector transfected *Entamoeba* trophozoites were resolved and probe with human anti-V1B. The *red* arrowheads show the predicated band corresponding to HepG2 and *E. histolytica* V1B subunits.

**Figure S6. V1B knockdown reduces macropinocytic cups**

**A**. *Entamoeba* trophozoites lysates (30 µg) from vector control (pTrigger without gene of interest) and V1B knockdown (KD) were resolved by SDS-PAGE followed by western blotting with human anti-V1B and anti-actin antibodies.

**B***.* Vector control and V1B KD trophozoites were incubated for 15 min at 37°C under steady state condition. Post incubation, trophozoites were probe with phalloidin and confocal images were acquired. The *white* arrowheads show pinocytic cups. Scale bars, 5 µm

**Supplementary Tables**

**Table S1.** Comparative sequence similarity of V-ATPase subunits from *E. histolytica*.

**Table S2.** List of primers used in current study.

**Table S3.** List of proteins detected in phosphatidic acid pulldown assays using mass spectrometry.

**Table S4**. Classification of cellular processes associated with proteins detected in phosphatidic acid pulldown assays.

**Supplementary Movie**

**Movie S1. Live cell imaging of GFP-V0d during macropinocytosis**

The movie illustrates the process of macropinocytosis, showing that GFP-V0d accumulates at regions of negative membrane curvature (the site of the “gulp”), increases at ruffling areas, and decline once the cup closes. GFP-V0d indicates in *green* channel and TRITC dextran in *red* channel. Scale Bar, 20 μm

**References**

Biller, L., J. Matthiesen, V. Kuhne, H. Lotter, G. Handal, T. Nozaki, Y. Saito-Nakano, M. Schumann, T. Roeder, E. Tannich, E. Krause, and I. Bruchhaus. 2014. The cell surface proteome of Entamoeba histolytica. *Mol Cell Proteomics*. 13:132-144.

Verma, K., and S. Datta. 2017. The Monomeric GTPase Rab35 Regulates Phagocytic Cup Formation and Phagosomal Maturation in Entamoeba histolytica. *J Biol Chem*. 292:4960-4975.
