## Supplementary material for "Evolutionary divergence V-ATPase function in macropinocytic cup remodeling": Table_S2.docx

Table S2: Gene-specific primer sequences of V-ATPase subunits used in the current study.

| **Putative V-ATPase subunit** | **Accession ID*** | **Primer Name** | **Primers sequence (5' to 3')** |
| --- | --- | --- | --- |
| V1A | EHI_043010 | V1A_SmaI_F | GCACCCGGGATGAACTTCGATACAGACAAAAAAG |
|  |  | V1A_XhoI_R | GACCTCGAGTTATTCTAAAAGAGTAGCAAAACGAGTGG |
| V1B | EHI_189850 | V1B_SmaI_F | GCACCCGGGATGGAAGCTTTCAAAATTACGCTGCTGC |
|  |  | V1B_XhoI_R | GACCTCGAGTTATAAGGTTCCTTCTTTGTCGTC |
| V1C | EHI_092600 | V1C_SmaI_F | GCACCCGGGATGACCCAAACTGGACTTTACC |
|  |  | V1C_XhoI_R | GACCTCGAGTTATCTTGAAACTAGTAACAACGGATTGAC |
| V0d | EHI_106350 | V0d_SmaI_F | GCACCCGGGATGCTTGGTGGATTATGGACATTCAATATGG |
|  |  | V0d_XhoI_R | GACCTCGAGTTAGAATAATCTAATAACGTTTTGATTG |

*The accession IDs of each V-ATPase subunits were recorded from Amoeba Informatics Resources (https://amoebadb.org/)
