## Supplementary figures and images for "Evolutionary divergence V-ATPase function in macropinocytic cup remodeling"

### S1.tif

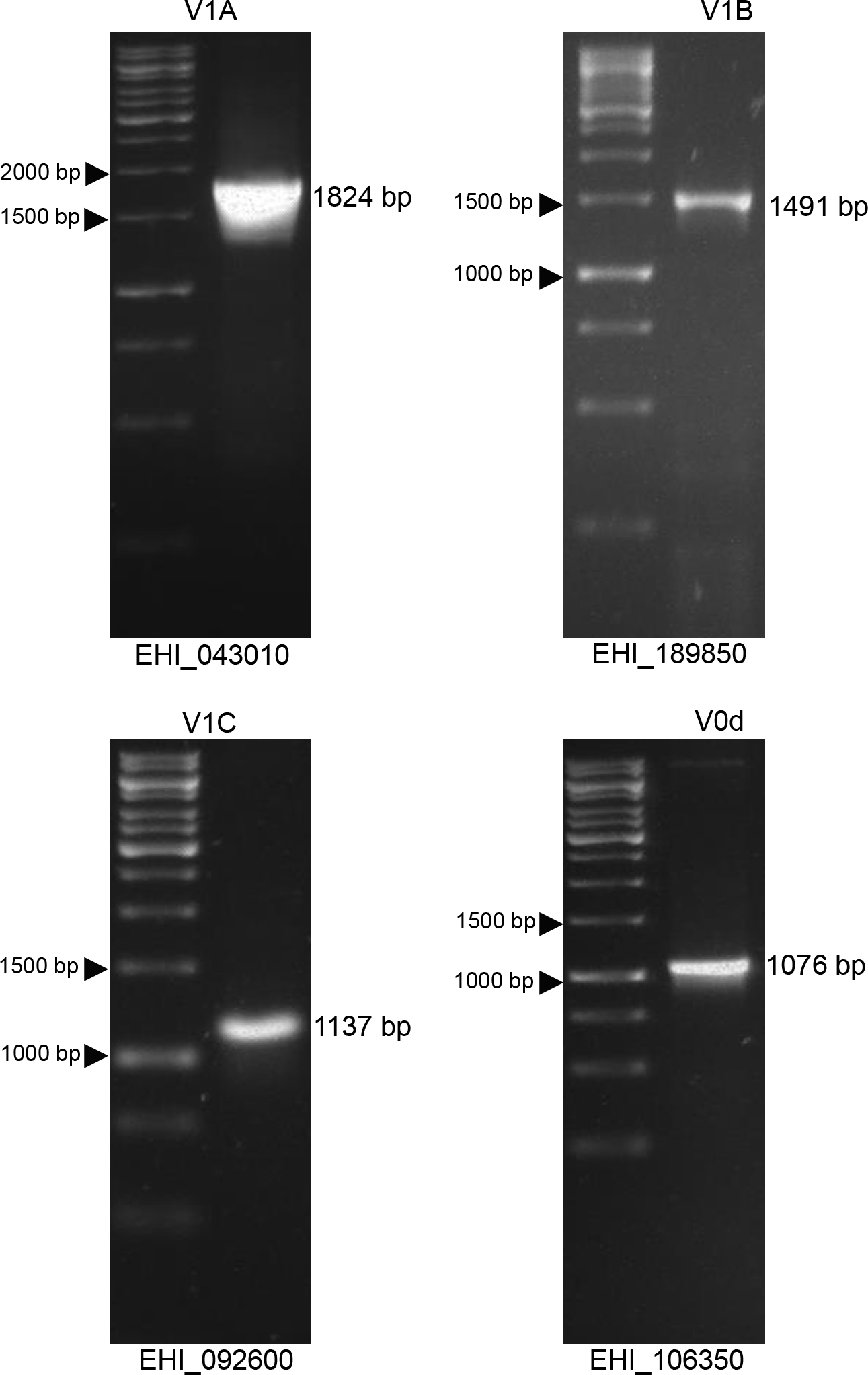

### S2.tif

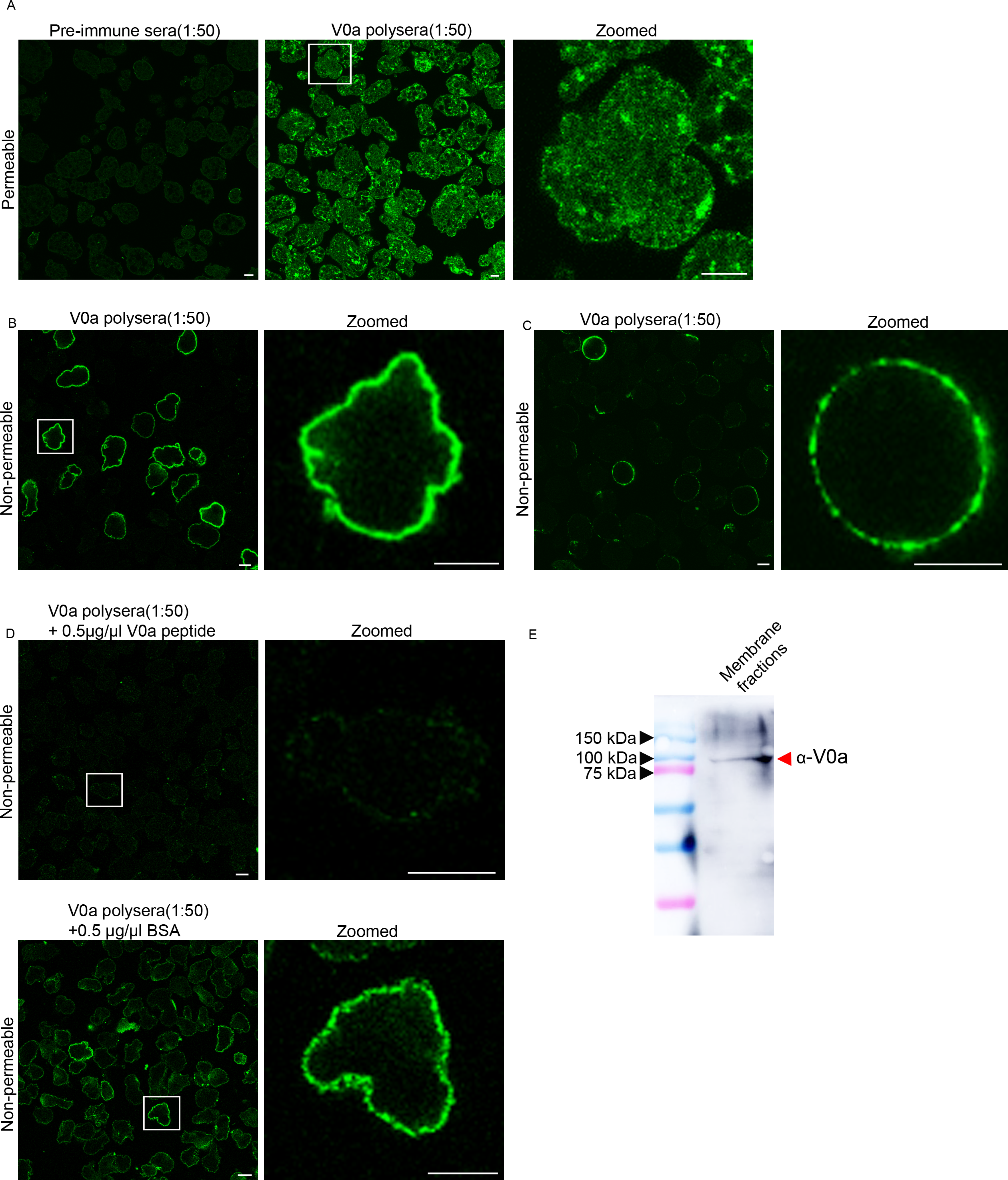

### S3.tif

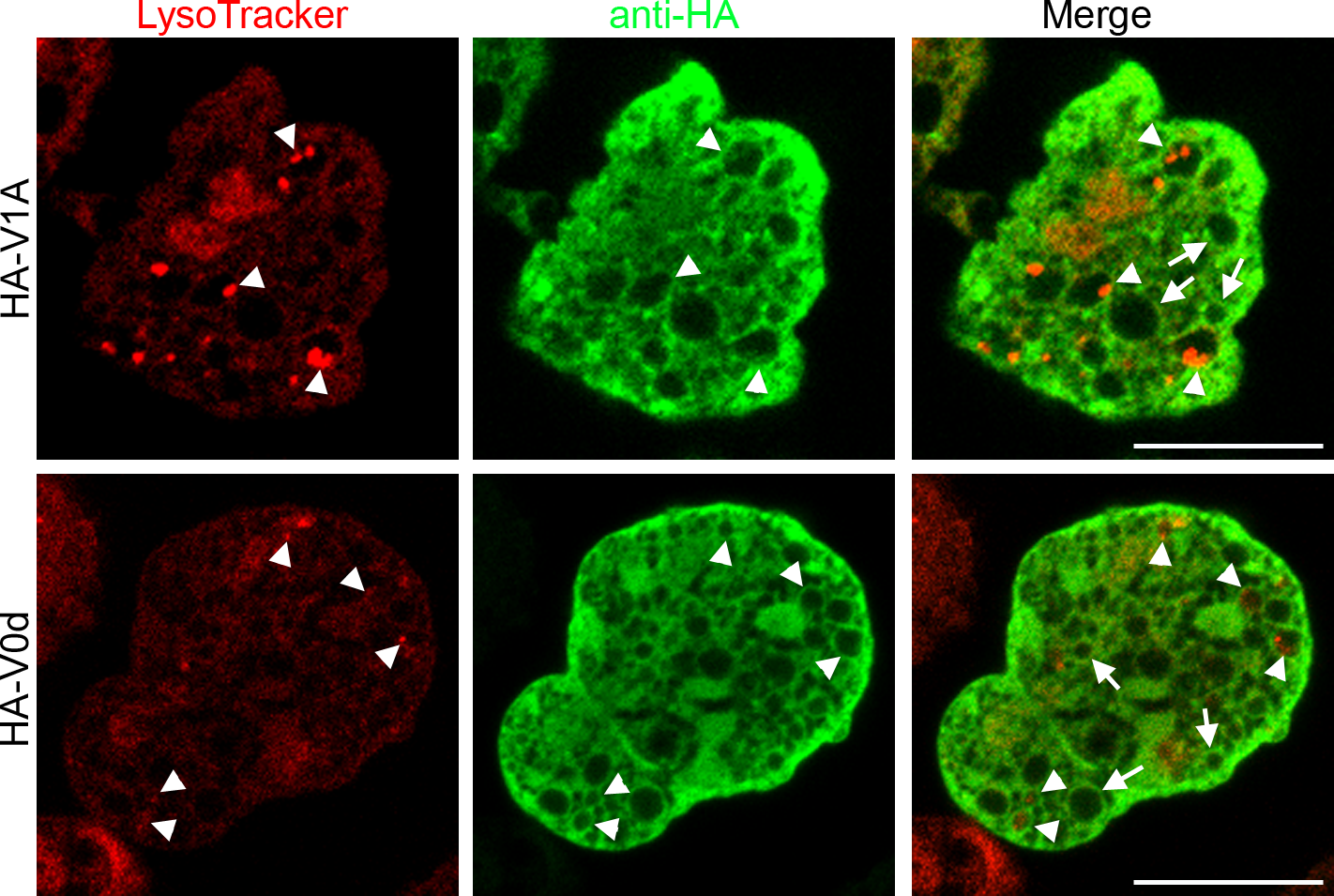

### S4.tif

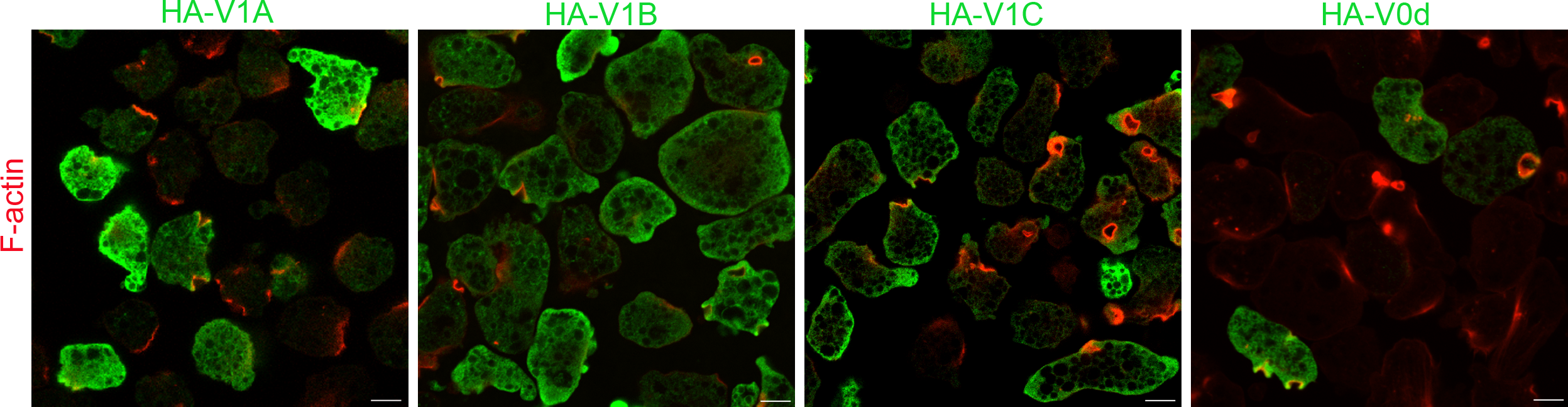

### S5.tif

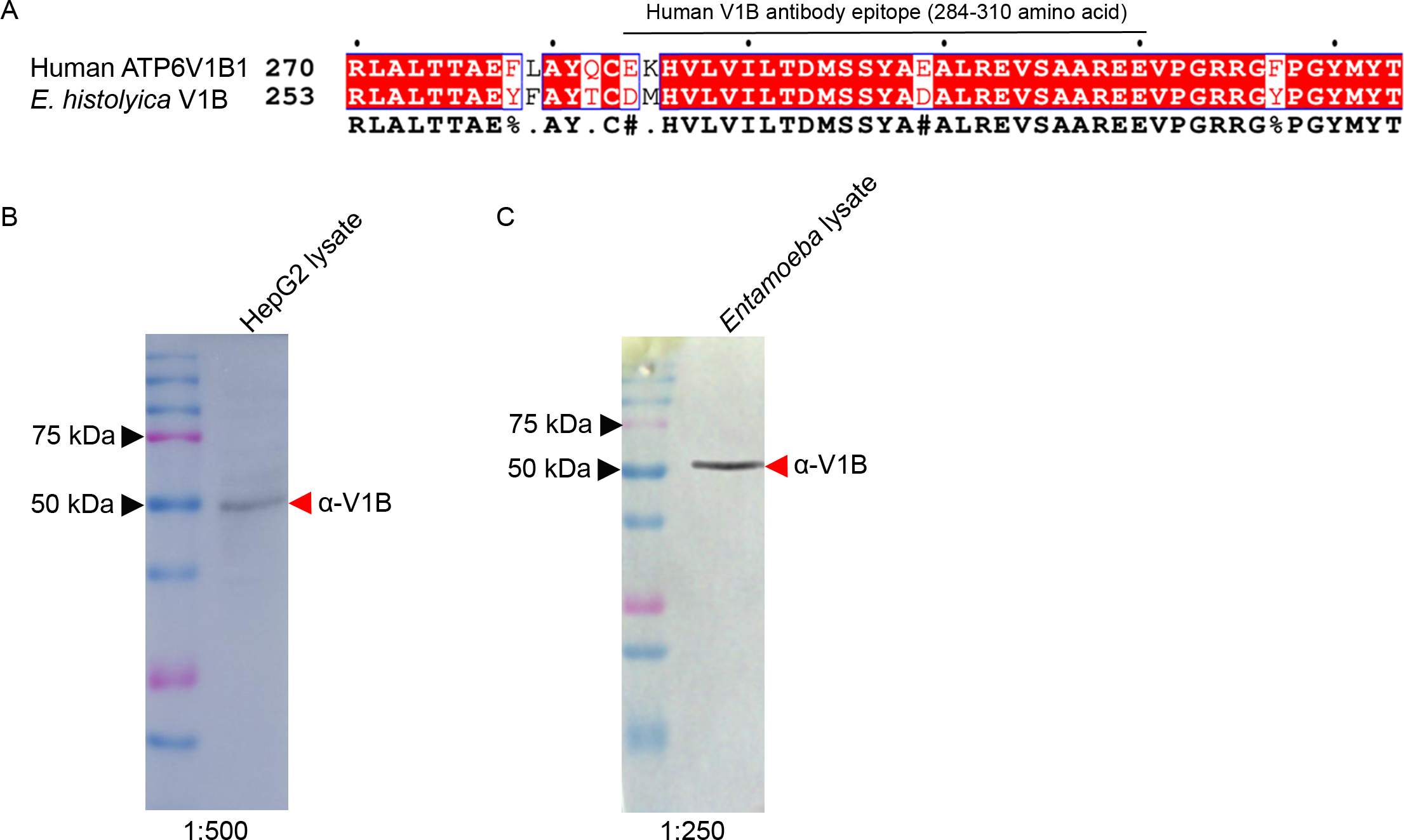

### S6.tif

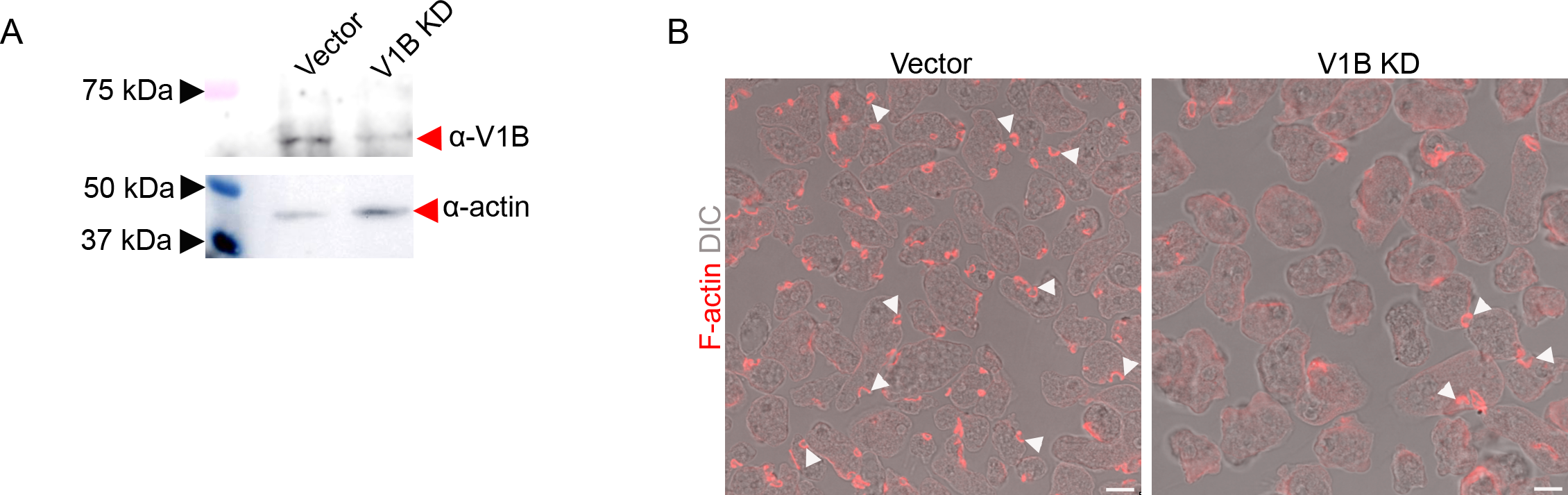
